# Single-cell RNA sequencing reveals functional heterogeneity and sex differences of glioma-associated brain macrophages

**DOI:** 10.1101/752949

**Authors:** Natalia Ochocka, Pawel Segit, Kacper Adam Walentynowicz, Kamil Wojnicki, Salwador Cyranowski, Julian Swatler, Jakub Mieczkowski, Bozena Kaminska

**Affiliations:** Laboratory of Molecular Neurobiology, Nencki Institute of Experimental Biology of the Polish Academy of Sciences, Warsaw, Poland; Postgraduate School of Molecular Medicine, Medical University of Warsaw, Warsaw, Poland; Laboratory of Cytometry, Nencki Institute of Experimental Biology of the Polish Academy of Sciences, Warsaw, Poland

## Abstract

Microglia are resident myeloid cells in the central nervous system (CNS) that control homeostasis and protect CNS from damage and infections. Microglia and peripheral myeloid cells accumulate and adapt tumor supporting roles in human glioblastomas that show prevalence in men. Cell heterogeneity and functional phenotypes of myeloid subpopulations in gliomas remain elusive. Single-cell RNA sequencing (scRNA-seq) of CD11b^+^ myeloid cells in naïve and GL261 glioma-bearing mice revealed distinct profiles of microglia, infiltrating monocytes/macrophages and CNS border-associated macrophages. We demonstrated an unforeseen molecular heterogeneity among myeloid cells in naïve and glioma-bearing brains, validated selected marker proteins and showed distinct spatial distribution of identified subsets in experimental gliomas. We found higher expression of MHCII encoding genes in glioma-activated male microglia, which was corroborated in bulk and scRNA-seq data from human diffuse gliomas. Sex-specific gene expression in glioma-activated microglia may be relevant to sex differences in incidence and outcomes of glioma patients.

## Introduction

Innate immune cells are abundant in the tumor microenvironment (TME) and play pivotal role in tumor progression and modulation of responses to therapy^1^. High number of macrophages within the TME have been associated with poor prognosis in many cancers, because those tumor-educated cells suppress antitumor immunity, stimulate angiogenesis and promote tumor invasion^2^. The central nervous system (CNS) is equipped with resident innate immune cells: microglia, and CNS border-associated macrophages (BAMs) that migrate to the CNS during the prenatal life and maintain a long-lasting population. In malignant gliomas, both local microglia and circulating monocytes migrate to the TME and differentiate into tumor supporting cells, commonly referred to as glioma-associated microglia and macrophages (GAMs). Reliable identification of specific subpopulations is hampered by a shortage of specific markers^3^. Transcriptome profiling of bulk CD11b^+^ cells isolated from human glioblastomas (GBMs) and rodent gliomas showed a mixture of protumorigenic and antitumorigenic phenotypes, and did not reveal consistent markers and pathways^4–6^. Recent reports showed that GAMs consist of diverse cell populations with likely distinct roles in tumor progression^7–10^. Dissecting the TME composition and functional heterogeneity of tumor-infiltrating immune cells would extend the understanding of glioma immune microenvironment and allow to modulate functions of distinct subpopulations for therapeutic benefits.

Sex differences in incidence (male-to-female ratio of 1.6:1), transcriptomes, and patient outcomes in adult GBM patients have been previously reported^11^. Sex-specific disease outcomes can be related to immune functions, because the efficacy of cancer immunotherapy in humans was shown to be largely depending on sex, with better outcomes in males^12^. In naïve mice, male microglia show enrichment of inflammation and antigen presentation-related genes, whereas female microglia have higher neuroprotective capacity^13,14^. Until now, sex differences have been largely unexplored in animal studies on glioma immunobiology.

Here, we used single-cell RNA sequencing (scRNA-seq) to decipher the composition and functions of GAMs in murine experimental GL261 gliomas grown in male and female mice. We demonstrate distinct transcriptional programs of microglia, monocytes/macrophages, and CNS BAMs. The identified microglia and monocyte/macrophage signature markers allow for a separation of these cells within glioma TME. Intracranial gliomas activate similar transcriptional networks in microglia and monocytes/macrophages present in TME. However, transcriptional responses of monocytes/macrophages are more pronounced and associated with activation of immunosuppressive genes. In males, microglia and a fraction of monocytes/macrophages infiltrating gliomas show higher expression of the MHCII genes suggesting stronger activation of male microglia. Altogether, this study demonstrates considerable cellular and functional heterogeneity of myeloid cells in TME and sex-specific differences in responses of myeloid cells to gliomas.

## Results

### Single-cell RNA-seq identifies myeloid cells with distinct expression profiles amongst CD11b^+^ cells from naïve and glioma-bearing brains

We employed a murine orthotopic GL261 glioma model, because tumors established from GL261 cells recapitulate many characteristics of human GBMs and are frequently used in studies of glioma immunology, immunotherapy, and in preclinical studies^15^. To assess the heterogeneity of GAMs in GL261 gliomas, we performed scRNA-seq on CD11b^+^ cells sorted from naïve and tumor-bearing brains of male and female mice (two replicates per group, two pooled mice per replicate) **(Figure 1a)**. We used naïve brains as controls, because scRNA-seq data for CD11b+ cells sorted from brains of naïve and sham-implanted animals did not indicate that the surgical procedure affects identified cell populations, their proportions and gene expression **(Supplementary figure 1e-h)**. The tumor-bearing animals were sacrificed 14 days post implantation. This time point corresponds to a pre-symptomatic stage of tumorigenesis, when GL261 tumors are restricted to a single hemisphere, show a substantial infiltration of peripheral monocytes/macrophages^16^, and no signs of necrosis that could affect perfusion **(Supplementary Figure 1a-d)**. Using fluorescence activated cell sorting, we sorted CD11b^+^ cells with a particularly high purity (>96%) and viability (~95%) **(Supplementary Figure 1d)**. We did not detect differences in the tumor size between sexes at this stage of tumor growth **(Supplementary Figure 1e)**.

**Fig. 1.**
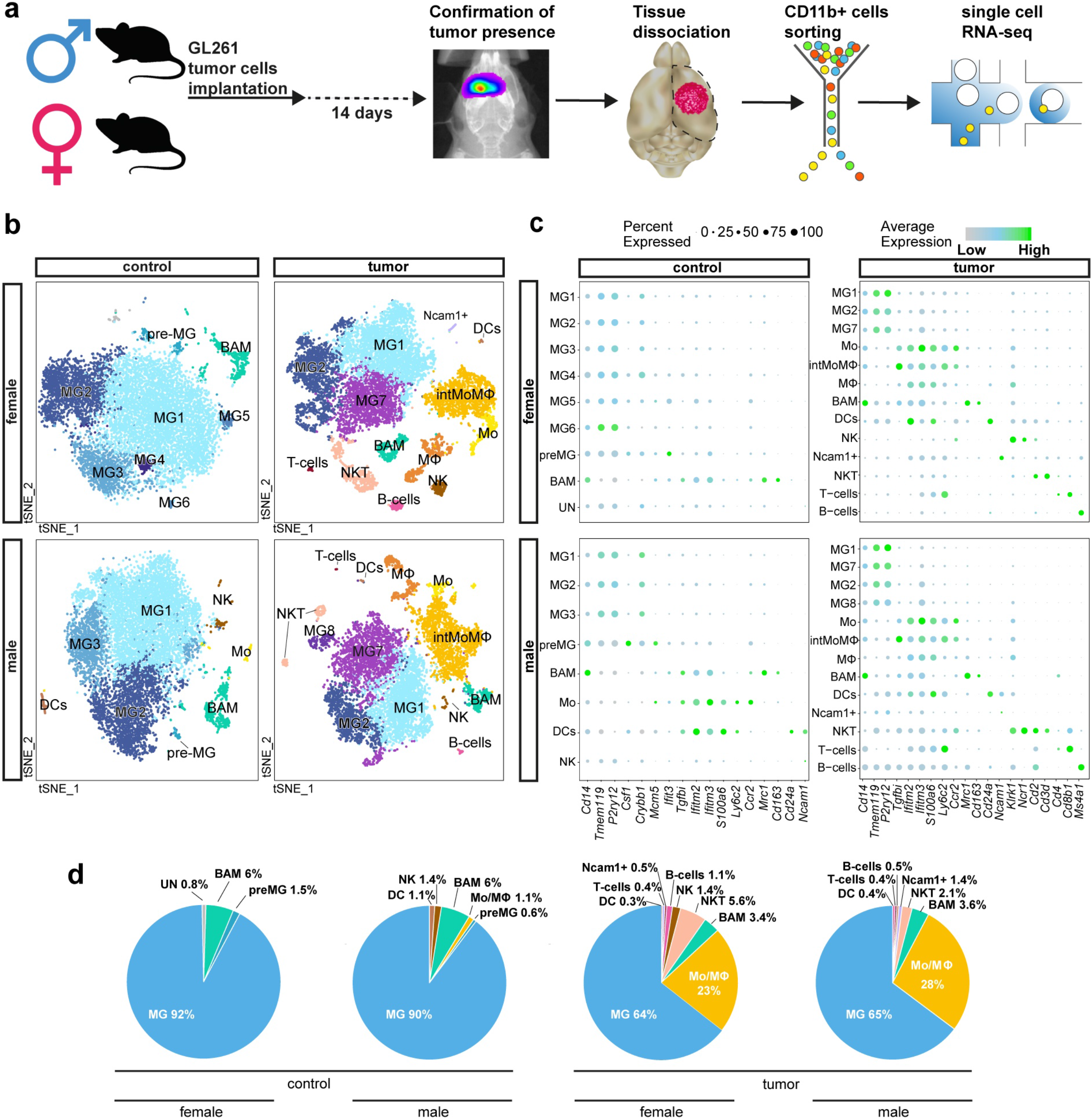
Identification of immune cell populations in control and tumor-bearing brain hemispheres. **a** Scheme of the experimental workflow. **b** t-SNE plot demonstrating clustering obtained for each group (female control, female tumor, male control, male tumor), 2 biological replicates were combined. Clusters annotations: MG – microglia, pre-MG – premature microglia, Mo – Monocytes, intMoMΦ – intermediate Monocyte-Macrophage, MΦ – macrophages, BAM – CNS border-associated macrophages, DCs – dendritic cells, Ncam1+− Ncam1 positive cells, NK – natural killer cells, NKT – natural killer T cells, B cells – B lymphocytes, T cells – T lymphocytes. **c** Expression of “signature” genes selected from the immune marker panel for identification of a cluster cell type (Supplementary Table 1). **d** Pie charts demonstrating distribution of the identified cell types across samples.

To resolve the molecular profiles of CD11b^+^ cells, we performed single-cell RNA sequencing. After quality control and adjusting for technical noise, single-cell transcriptomic profiles for 40,401 cells and 14,618 genes were selected for the analysis (see Methods). We visually inspected the transcriptomic diversity of computed clusters, projecting the data onto two dimensions by t-distributed stochastic neighbor embedding (t-SNE) **(Figure 1b)**. To characterize the cell identity of the obtained clusters, we applied the immune cell marker panel **(Figure 1c)** created with the literature-based markers **(Supplementary Table 1)**^3,7,8,10,17–30^. The cell identities were inferred by identifying significantly overexpressed genes in each cluster.

Unsupervised clustering of each group demonstrated a similar number of clusters between sexes **(Figure 2b)**. To avoid over-fitting, we used the same parameters for all cluster analyses (see Methods) and this might have resulted in different stratification of the analyzed conditions. However, this analysis was done primarily to select cells for further processing.

**Fig. 2.**
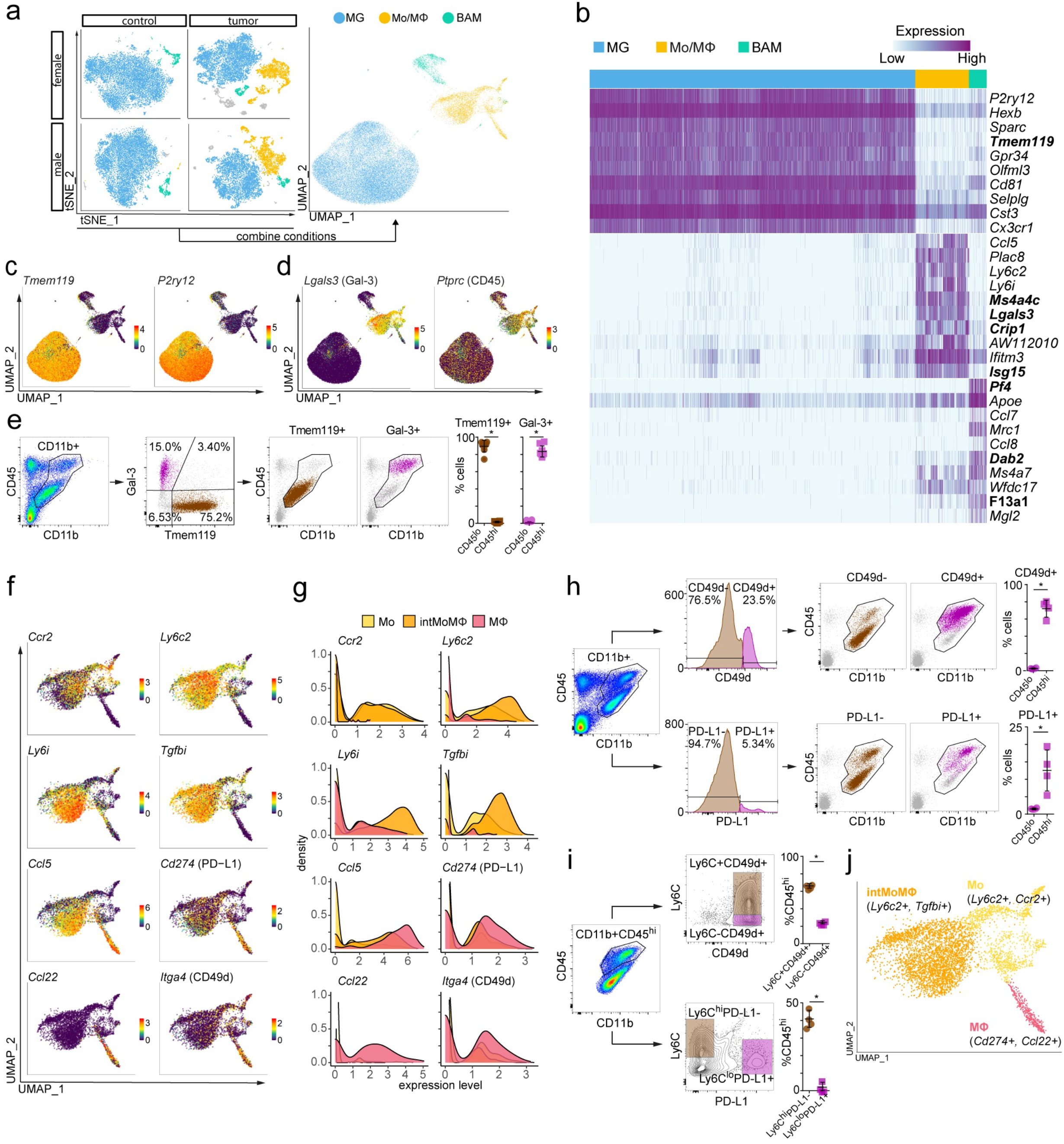
Transcriptomic characterization of main myeloid subpopulations. **a** Projection of cells combined from clusters identified as microglia, monocytes/macrophages (Mo/MΦ), and BAMs from all groups. **b** Top 10 differentially expressed genes for the three main identified cell populations, new marker candidates are in bold. **c-d** Feature plots depicting genes highly expressed in MG (**c**) and MoMΦ (**d**). **e** Flow cytometric analysis of the distribution of Tmem119 and Gal-3 protein markers within CD11b+ cells and projection of Tmem119+ and Gal-3+ cells onto CD45/CD11b graphs, dot plots demonstrate percentages of Tmem119+ and Gal-3+ cells within CD45^hi^ and CD45^lo^ groups (n=8, 4 males and 4 females, Mann-Whitney U test, *<0.05). **f** Feature plots depicting distribution of the expression of genes discriminating monocytes/macrophages (Mo/MΦ), Monocytes (Mo), Monocyte-Macrophage intermediate (intMoMΦ), and Macrophage (MΦ) subpopulations. **g** Density plots demonstrating the expression level of markers discriminating Mo/MΦ subpopulations. **h** Flow cytometric analysis of CD49d and PD-L1 proteins within CD11b+ cells and their projection onto CD11b/CD45 graphs, dot plots demonstrate percentages of CD49d+ and PD-L1+ cells within CD45^hi^ and CD45^lo^ groups (n=4, 2 males and 2 females; Mann-Whitney U test, *<0.05). **i** Flow cytometric analysis of the distribution of the markers discriminating Mo/MΦ subpopulations within CD11b+CD45^hi^ cells, dot plots demonstrate percentage of CD11b+CD45^hi^ cells that belong to the defined populations (n=4, 2 males and 2 females; Mann-Whitney U test, *<0.05). **j** UMAP plot shows clusters of Mo/MΦ subpopulations.

For naïve female and male CD11b^+^ cells, 9 and 8 clusters were obtained, respectively. Gene expression profiles underlying a specific cluster could reflect different functions of contained cells or their different origin from various brain structures. To dissect functional meaning of gene expression underlying microglial clusters (MG), we explored the available information on microglial phenotypes^20,21,31,34^. The predominant cluster MG1 characterized by relatively high expression of microglia enriched genes (*Crybb1, Cst3, P2ry12*, *Pros1*)^31^ may reflect a subpopulation of homeostatic microglia. Cluster MG2 is characterized by a high expression of immediate early genes (*Jun, Junb, Jund, Fos, Egr1, Klf6, Aft3*) encoding transcription factors, and may encompass a subpopulation of transcriptionally active cells **(Supplementary Table 5)**. MG2 shows also an increased expression of *Nfkbia* encoding an NFκB inhibitor alpha. Cluster MG3 is marked by high expression of genes coding for signaling inhibitor: *Bmp2k* and transcriptional repressors: *Bhlhe41*, *Ncoa3* and *Notch2* **(Supplementary Table 5)**. The transcription factors Bhlhe40 and Bhlhe41 directly repress the expression of lineage-inappropriate genes in alveolar macrophages^32^. *Ncoa3* in association with nuclear receptors represses expression of inflammation mediators and activates genes encoding anti-inflammatory mediators^33^. Three microglial clusters represented by a smaller cell number: MG4, MG5 and MG6 were identified in female naïve CD11b^+^ cells. MG4 did not show a cluster-specific genes. MG5 showed increased expression of myelin specific genes: *Plp1*, *Pltp* and *Mbp*, which are found in microglia with increased myelin uptake^34^. Among top highly expressed genes in MG6, we found *Cd63* and *Cd9* encoding proteins that are enriched in extracellular vesicles^35^.

In both male and female CD11b^+^ cells from naïve brains we identified Pre-MG cluster characterized by an increased expression of microglial genes (*Tmem119*, *P2ry12, Crybb1*) and genes characteristic for their premature state (*Csf1*, *Mcm5*, *Ifit3*)^21^ **(Figure 1c)**. Pre-MG upregulated genes encoding a cysteine protease inhibitor (*Cst7)*, cytokines (*Mif* and *Csf1)*, chemokines (*Ccl12*, *Ccl3*, *Ccl4*), genes involved in a response to interferon (*Ifit1*, *Ifit3*, *Ifit3b*, *Ifitm3*, *Irf7*), and genes implicated in a ubiquitin-like process of ISG-ylation (*Isg15*, *Usp18*) that are activated during inflammation **(Supplementary Table 5)**. These microglia could represent surveying cells fitted to rapidly respond to homeostasis dysfunction.

Among CD11b^+^ cells from tumor-bearing hemispheres we identified 13 clusters for both sexes. In tumor infiltrating microglia, besides the presence of previously described clusters MG1-2, we found an MG7 cluster characterized by increased expression of genes encoding the components of major histocompatibility complex (MHC) class I (*B2m, H2-D1*, *H2-K1*) and MHC class II (*H2-Oa*, *H2-DMa)*, *Bst2 and Lgals3bp*, upregulation of which has been reported in disease-associated microglia^36,37^, and *Ccl12* encoding a cytokine critical for CCR2^+^ monocytes recruitment^38^. Among tumor-infiltrating male microglia, we found cluster MG8 characterized by a high expression of genes encoding proliferation-related proteins (*Stmn1, Tubb5, Tuba1b, Cdk1*, *Top2a*), which is consistent with the observed proliferation of glioma-activated microglia, as previously reported^6,39^. Microglia from tumor animals show also upregulation of *Timp2, Serpine2, Cst7* and *Ctsd*, genes encoding proteases or their modulators participating in reorganization of extracellular matrix, which may reflect invasion supporting properties. The identified MG clusters may represent a transient, intermediate activation states of microglia.

In naïve brains, microglia (MG) comprised the vast majority of all sorted cells (91% in females, 90% in males), whereas BAMs constituted 6% of cells in both sexes **(Figure 1d)**. Amongst CD11b^+^ cells from male controls, we found a small subset of monocytes (Mo, *Ly6c2*^+^, *Ccr2*^+^), natural killer (NK, *Ncam1*^+^), and dendritic cells (DC, *Cd24a*^+^).

In tumor-bearing brains, microglia were still the most abundant cell population (64% in females, 65% in males), although their proportion decreased due to infiltration of monocytes/macrophages (Mo/MΦ), forming the second main myeloid cell population of the TME (23% in females, 28% in males) **(Figure 1d)**. For both sexes, we identified 3 clusters of infiltrating Mo/MΦ that could be further characterized by an inflammatory monocyte signature – Mo (*Ly6c2*^hi^, *Ccr2*^hi^, *Tgfbi*^lo^), an intermediate state of monocyte and macrophage signature – intMoMΦ (*Ly6c2^hi^*, *Tgfbi*^hi^), and a differentiated macrophage signature – MΦ (*Ly6c2*^lo^, *Ifitm2*^hi^, *Ifitm3*^hi^, *S100a6*^hi^) **(Figure 1c)**. These results demonstrate dynamic changes in monocytes/macrophages infiltrating gliomas. We found minor populations of NK cells, DCs, natural killer T cells (NKT), and a marginal fraction of B and T cells. CD11b^+^ is not expressed on lymphocytes, but rare CD11b+ lymphocytes (<1%) may appear after activation of the immune response^40,41^. Nevertheless, a vast majority of cells were MG, Mo/MΦ, and BAM.

### Assessment of new and known cell type specific markers

To identify the molecular features that distinguish naïve and tumor-associated myeloid cells, we performed further analyses on major cell subpopulations. From all the conditions and replicates, we extracted only the cells identified as microglia (MG), monocytes/macrophages (Mo/MΦ), and BAMs. In all conditions, both scRNA-seq replicates similarly contributed to these results **(Supplementary Figure 2)**, demonstrating good reproducibility. The combined three cell subpopulations, projected on the two-dimensional space using a Uniform Manifold Approximation and Projection (UMAP) algorithm, formed three separate groups **(Figure 2a**, **Supplementary Figure 3)**. This observation demonstrates a predominance of a biological signal over technical artifacts or batch effects. To confirm cell identities, we performed differential expression analyses between three subpopulations of CD11b^+^ cells. Among the most highly upregulated genes in each group (see Methods for details of differential gene expression analysis), we found the well-known microglial genes – *P2ry12*, *Sparc*, *Tmem119*, *Gpr34*, *Selplg*, *Cx3cr1*^19,42^ in MG, monocyte – *Ly6i*, *Ly6c2*, and macrophage genes – *Ifitm3*^10^ in Mo/MΦ, and BAM genes – *Apoe*, *Ms4a7*, *Mrc1*^43^ in BAMs **(Figure 2b)**. The expression of *Tmem119*, *Cx3cr1*, *P2ry12*, *Gpr34*, *Olfml3*, and *Sparc* was enriched only in microglia **(Figure 2b**, **Supplementary Figure 5a)**. Other genes expressed at a high level in microglia were also highly expressed in BAMs (*Cd81*), BAMs and Mo/MΦ (*Hexb, Cst3*) or were found only in a fraction of cells (*P2ry13* gene was expressed by less than 75% of MG cells) **(Supplementary Figure 5a)**. For Mo/MΦ, we found enriched expression of previously reported genes such as *Ifitm2*, *S100a6*, and *S100a11*^10^, as well as novel genes, namely *Ms4a4c*, *Lgals3*, *Crip1* and *Isg15* **(Figure 2b**, **Supplementary Figure 5b)**. *Ifitm3* was highly expressed by the Mo/MΦ population, but appeared in a substantial fraction of MG, showing its low specificity in monocytes/macrophages within glioma TME. Highly expressed genes in BAMs were *Apoe* and *Ms4a7*, recently proposed as markers of CNS border macrophages^43^. However, we found these genes also highly expressed by Mo/MΦ, suggesting that *Apoe* and *Ms4a7* are not exclusive for BAMs in TME. *Mrc1* showed high expression restricted to BAMs. Additionally, we found *Pf4*, *Dab2 and F13a1* highly and specifically expressed by BAMs **(Figure 2b**, **Supplementary Figure 5c)**.

We aimed to identify markers for separation of microglia and macrophages in TME. From the top differentially expressed genes (ranked by the average log fold-change value, all with adjusted (Bonferroni correction) p-value < 10^−100^) in the MG and Mo/MΦ groups **(Figure 2b)**, we selected candidate genes with enriched expression in a majority of cells in the group of interest – *Tmem119* (MG) and *Lgals3* (Mo/MΦ) **(Figure 2c,d)**. *Tmem119* was proposed as a microglia marker by Bennet *et al.* (2017)^19^, who confirmed its utility in CNS inflammation and nerve injury. *Lgals3* encodes galectin-3 (Gal-3), a lectin involved in tumor immunosuppression^44^. Gal-3 is produced and secreted by macrophages, regulates IL-4 induced alternative macrophage activation^45^ and acts as monocyte/macrophage chemoattractant.

We assessed Tmem119 and Gal-3 expression in CD11b^+^ cells from tumor-bearing hemispheres by flow cytometry at day 14 post-implantation **(Figure 2e)**. Brains were mechanically processed and dissociated enzymatically with DNase I to preserve a Tmem119 surface marker (see **Supplementary Figure 6a** and Methods). Gal-3 and Tmem119 allowed for the discrimination of two populations: Tmem119^+^Gal-3^−^ (75.2% of cells) and Tmem119^−^ Gal-3^+^ (15.0% of cells), whereas Tmem119^+^Gal-3^+^ population was minor **(Figure 2e)**. These results correspond to scRNA-seq analysis in which 60% and 21.9% of CD11b+ cells expressed only *Tmem119* or *Lgals3*, respectively. We assessed Tmem119 and Gal-3 expression in CD11b^+^CD45^lo^ and CD11b^+^CD45^hi^ cells in order to compare these marker candidates with the previously used method. Interestingly, the two methods produced similar separation, as 89.5% of CD11b^+^CD45^lo^ cells were Tmem119^+^ and 83.4% of CD11b^+^CD45^hi^ cells were Gal-3^+^ **(Figure 2e)**.

Among the highly upregulated Mo/MΦ genes, we found candidates enriched in discrete subpopulations of Mo/MΦ. The high *Ly6c2* expression was found in a large cell fraction, which could be further divided into *Ly6c2*^hi^*Ccr2*^hi^ monocytes (Mo) and *Ly6c2*^hi^*Tgfbi*^hi^ monocyte/macrophage intermediate cells (intMoMΦ) **(Figure 2f,g)**. The remaining cells resembled differentiated tissue macrophages (MΦ), because they lacked the markers of the cytotoxic monocytes (*Ly6c2*, *Ccr2)* and had a strong “macrophage signature” (*Ifitm2*^hi^, *S100a6*^hi^, *S100a11*^hi^) **(Supplementary Figure 5b)**.

Notably, we found a population of MΦ expressing *Ccl22* and *Ccl5* genes, encoding chemokines important for T-cell recruitment^46,47^ and *Cd274*, a gene encoding an immune checkpoint protein PD-L1 **(Figure 2f,g)**. Such expression pattern suggests a putative role of these cells in mediating the immunosuppressive response. Flow cytometric analysis confirmed that PD-L1 expression is restricted to CD11b^+^CD45^hi^ population **(Figure 2h)**. Distribution of Ly6C and PD-L1 among CD45^hi^ population indicates that those proteins denote distinct populations among peripheral myeloid cells infiltrating gliomas: Ly6C^hi^PD-L1^−^ intermediate monocyte/macrophages (intMoMΦ) and Ly6C^lo^PD-L1^+^ differentiated macrophages (MΦ) **(Figure 2i**, **Supplementary Figure 6)**. Thus, we identified genes enriched in the monocyte/macrophage subpopulations **(Figure 2j)**.

We also examined the expression of genes recently proposed as specific markers of monocytes/macrophages in gliomas: *Itga4*^7^, *Hp*, *Emilin2*, *Sell*, and *Gda*^23^ **(Figure 2f**, **Supplementary Figure 5d)**. The expression of *Itga4* (CD49d) was low and limited mostly to the MΦ subpopulation, resembling differentiated macrophages and expressing a high level of *Cd274* (PD-L1). However, flow cytometric analysis showed that the CD49d protein is expressed by 72.1% of CD11b^+^CD45^hi^ cells **(Figure 2h)**, out of which 66.2% are Ly6C^hi^ and 24.5% Ly6C^lo^ **(Figure 2i)**, demonstrating that CD49d protein is expressed in both monocytic and macrophage fraction of bone marrow-derived macrophages (BMDM). CD49d was not found in CD11b^+^CD45^lo^ cells, which corroborated its specificity towards monocyte/macrophage compartment. The expression of *Hp*, *Emilin2*, *Sell*, *Gda*, the markers suggested in recent meta-analysis of bulk RNA-seq data sets and validated at RNA and protein levels^23^, was found in the fraction of Mo (Ly6c*2*^hi^, Ccr2^hi^) **(Supplementary Figure 5e)**. *Tgm2* and *Gpnmb*, previously reported as the genes commonly upregulated by GAMs across different glioma animal models and in a bulk RNA-seq of patient-derived samples^3^ was limited to the small fraction of Mo/MΦ **(Supplementary Figure 5e)**. This observation shows how bulk RNA-seq results may be biased by genes expressed at a high level in a small subset of cells.

Summarizing, we validated the expression of known markers at the single-cell level and obtained a coherence of selected microglia and BAM markers in our data set with literature data. In contrast, Mo/MΦ in TME showed substantial heterogeneity that is likely related to their differentiation state.

### Distinct gene expression profiles of glioma-associated microglia and monocytes/macrophages

Distribution of cells according to the experimental conditions (naïve versus tumor) revealed separation of functional subgroups of microglia. This separation was further supported by the unsupervised clustering that led to clusters either highly enriched in the cells from naïve brains representing *homeostatic microglia* (Hom-MG) or clusters dominated by cells originating from the tumor-bearing hemispheres representing *glioma-activated microglia* (Act-MG) **(Figure 3a**, **Supplementary Figure 7)**. This result demonstrates activation of microglia within TME. Mo/MΦ cell fraction is composed of TME infiltrating monocytes/macrophages. BAMs from naïve and tumor-bearing brains distributed evenly and did not show any clusters of cells originating predominantly from tumor-bearing hemispheres **(Figure 3a)**.

**Fig. 3.**
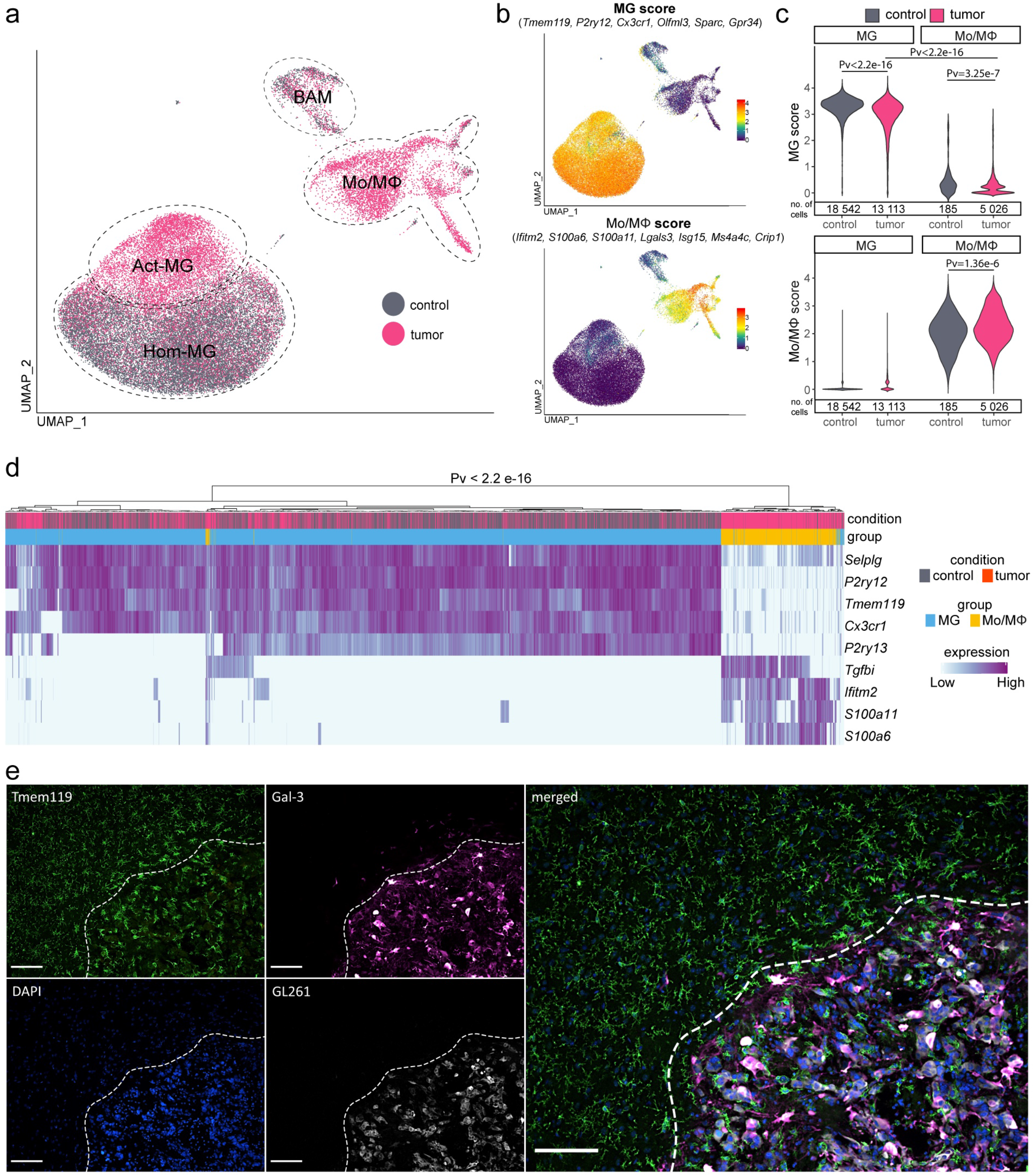
Tumor-derived microglia and macrophages form separate cell populations. **a** UMAP plots demonstrate the distribution of CD11b+ cells from naïve and tumor-bearing mice. **b** Distribution of MG and Mo/MΦ “signature” gene scores (presented as an average of expression of the selected genes). **c** Density plots of MG and Mo/MΦ scores across MG and Mo/MΦ populations demonstrating no overlap of a specific “signature” between the two cell populations, Wilcoxon signed-rank test. **d** Cell hierarchical clustering according to the expression of reported macrophage markers demonstrating bimodal cell distribution, Fisher’s exact test. **e** Immunohistochemical staining for microglia (Tmem119+, Gal-3-) and Mo/MΦ (Tmem119-, Gal-3+) shows the localization of specific immune cells within the tumor and its surroundings in female animal (for male see Supplementary Figure 8); a dashed line marks the tumor edge; scale −100 μm.

Using MG and Mo/MΦ scores (defined as an average of expression levels of genes restricted to and highly expressed in a given population) **(Figure 3b)**, we examined whether microglia and macrophage “signature” gene expression is modified in TME. We found a shift towards the lower “microglia signature” score in MG from the tumor-bearing brains compared to those from the naïve brains **(Figure 3c)**. Still, the “microglia signature” in MG from the tumor was strong and distinguishable from Mo/MΦ, allowing for clear separation of the two cell populations. Similarly, the “macrophage signature” score was high and distinctive for the Mo/MΦ population. Using selected markers, we performed hierarchical clustering of cells according to the expression of reported microglia and macrophage markers, resulting in clear separation of microglia and Mo/MΦ **(Figure 3d)**. This observation indicated that the expression of signature genes is retained even under the strong influence of the glioma microenvironment.

We separated distinct CD11b+ subpopulations: microglia (Tmem119^hi^) and monocyte/macrophages (Gal-3^hi^) by flow cytometry **(Figure 2e)**. Using immunostaining we studied spatial localization of those populations in TME. We demonstrate that Tmem119^+^ microglia adopt an amoeboid morphology in the tumor proximity and localize abundantly at the tumor edge, whereas Gal-3^+^ macrophages accumulate mostly within the tumor mass in both female **(Figure 3e)** and male animals **(Supplementary Figure 8)**. This finding confirms previous reports demonstrating that microglia occupy the tumor periphery and monocytes/macrophages localize mostly within the tumor core^9,48^.

Altogether, we show that microglia undergo glioma-induced activation associated with slight reduction of “microglia signature” gene expression, but both microglia and monocytes/macrophages retain expression of “signature” genes within TME. Staining for Tmem119 and Gal-3 separates microglia and monocytes/macrophages in murine gliomas.

### Transcriptional networks induced in microglia by glioma are present and more pronounced in infiltrating monocytes/macrophages

As demonstrated above, microglia and monocytes/macrophages have distinct gene expression profiles. To elucidate their roles in supporting glioma growth, we examined the transcriptional networks activated in MG and Mo/MΦ in TME. Firstly, we extracted genes highly upregulated in microglial cells from glioma-bearing brains (significantly upregulated genes in Act-MG compared to Hom-MG). Subsequently, we compared those profiles in Act-MG and Mo/MΦ cells **(Figure 4a)** to find genes either common or specific for each subpopulation. We found that the majority of genes upregulated in the Act-MG are also expressed by Mo/MΦ, and their expression is usually higher in Mo/MΦ than Act-MG **(Figure 4b,c)**. Among commonly induced genes, we found *Ifitm3* and a group of genes encoding MHCII proteins (*H2-Aa*, *H2-*Ab1, *H2-D1*, *H2-K1*). Expression of *Ifitm3* has been reported to demarcate macrophages from microglia^10^. We demonstrate that *Ifitm3* is highly expressed in monocytes/macrophages, but also in glioma-activated microglia (Act-MG) **(Figure 4b,c)**. Act-MG showed a high expression of *Ccl3*, *Ccl4*, and *Ccl12* (chemokine-encoding genes) when compared to Mo/MΦ. In contrast, Mo/MΦ were characterized by high expression of *Ifitm2* and *Ccl5* genes **(Figure 4b,c)**.

**Fig. 4.**
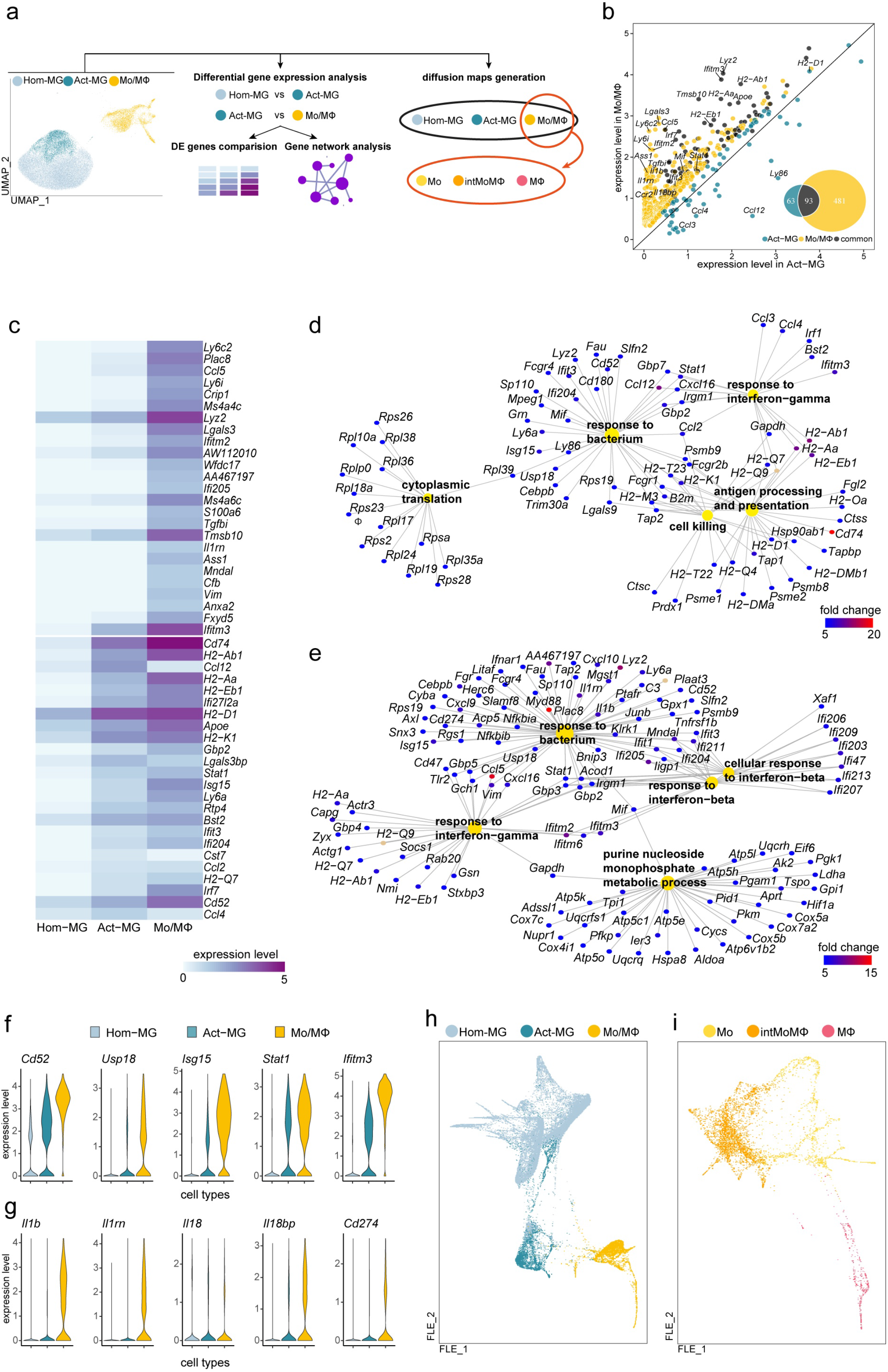
Functional analysis of glioma-activated microglia in comparison to tumor-infiltrating monocytes/macrophages. **a** Scheme of the analytical approach. **b** Scatter plot depicting expression level of differentially upregulated genes in Act-MG and Mo/MΦ. **c** Heatmap showing the comparison of expression of top 25 upregulated genes in Hom-MG vs Act-MG and Act-MG vs Mo/MΦ. **d-e** Gene Ontology analysis of biological processes for genes upregulated in (**d**) Act-MG compared to Hom-MG and (**d**) Mo/MΦ compared to the Act-MG. **f-g** Expression level of selected genes expressed specifically in distinct subpopulations **h-i** Visualization of cells projection on two dimensional FLE (force-directed layout embedding) space.

Next, we performed Gene Ontology (GO) analysis of biological processes on two sets of genes – genes significantly upregulated in Act-MG compared to Hom-MG **(Figure 4d)** and genes significantly upregulated in Mo/MΦ compared to the Act-MG **(Figure 4e)**. Gene expression in Act-MG was enriched in terms “cytoplasmic translation”, whereas terms “purine monophosphate metabolic process” were enriched in Mo/MΦ. All other terms were directly related to the immune function and largely shared between upregulated genes in MG and Mo/MΦ. Both populations showed induction of genes related to “response to bacterium” and “response to interferon-gamma”; however, those terms encompassed the broader number of genes for Mo/MΦ. In addition, Mo/MΦ demonstrated the enrichment of “response to interferon-beta” genes. The genes coding for MHCII components (e.g. *H2-Aa*, *H2-Ab1*, *H2-Eb1*) are upregulated in both MG and Mo/MΦ, however, in MG these genes are represented under “antigen processing and presentation” and “response to interferon gamma” terms and in Mo/MΦ they are represented only under “response to interferon gamma” term.

Several shared genes (*Cd52, Stat1, Isg15*, *and Usp18)* were expressed at a higher level in Mo/MΦ compared to their levels in Act-MG **(Figure 4f)**. Proteins encoded by those genes are involved in immune responses: CD52 mediates co-stimulatory signals for T-cell activation and proliferation^49^; Stat1 is a mediator of interferon signaling; Isg15 stabilizes Stat1 preventing premature termination of an inflammatory response^50^; Usp18 negatively regulates *Stat1* expression and termination of interferon-induced genes^51^. Such expression patterns may indicate that both microglia and monocytes/macrophages initiate some elements of the immune response, with more prominent activation in monocytes/macrophages. Among genes that were highly expressed in Mo/MΦ, we found *Il1b* coding for an inflammatory cytokine IL-1β along with *Il1rn* and *Il18b* coding for the inhibitors of pro-inflammatory cytokines **(Figure 4g)**. These data, together with the high expression of *Cd274* coding for PD-L1 in Mo/MΦ, suggest stronger activation of immunosuppressive pathways in monocytes/macrophages **(Figure 4g)**.

Additionally, we used scSVA tool^52^ to generate single-cell diffusion maps and to obtain visualization of our dataset on force-directed layout embedding (FLE) **(Figure 4h,i)**. Using previously assigned cell identity labels, we demonstrate with a different computational approach, how cells from each cluster are projected onto two-dimensional space. Analysis showed similar patterns of cell distribution for Hom-MG, Act-MG and Mo/MΦ cells on FLE **(Figure 4h)** as on UMAP **(Figure 3a)**, as well as for subpopulations of Mo/MΦ cells (**Figure 4i** and **Figure 2j**, respectively). Expression profiles of monocyte and macrophages suggest to some extent that transition from monocytes to intermediate monocyte-macrophages to differentiated macrophages takes place in glioma microenvironment.

### Sex-related differences in microglial expression of MHCII genes

Sex is an important prognostic marker in GBM patients influencing incidence and disease outcomes^11^. Differences between male and female microglia in naïve mice have been reported^13,14^. We examined whether there are sex-related differences in gene expression in main myeloid populations in gliomas. The unsupervised cell clustering showed that microglia from glioma-bearing brains, but not from naïve brains, segregate into clusters that are enriched either in cells originating from female or male **(Figure 5a**, **Supplementary Figure 10a)**. Similarly, we observed the sex-driven cell grouping within the intMoMΦ subpopulation, pointing to differences in immune cell activation in male and female mice **(Figure 5a**, **Supplementary Figure 10b)**.

**Fig. 5.**
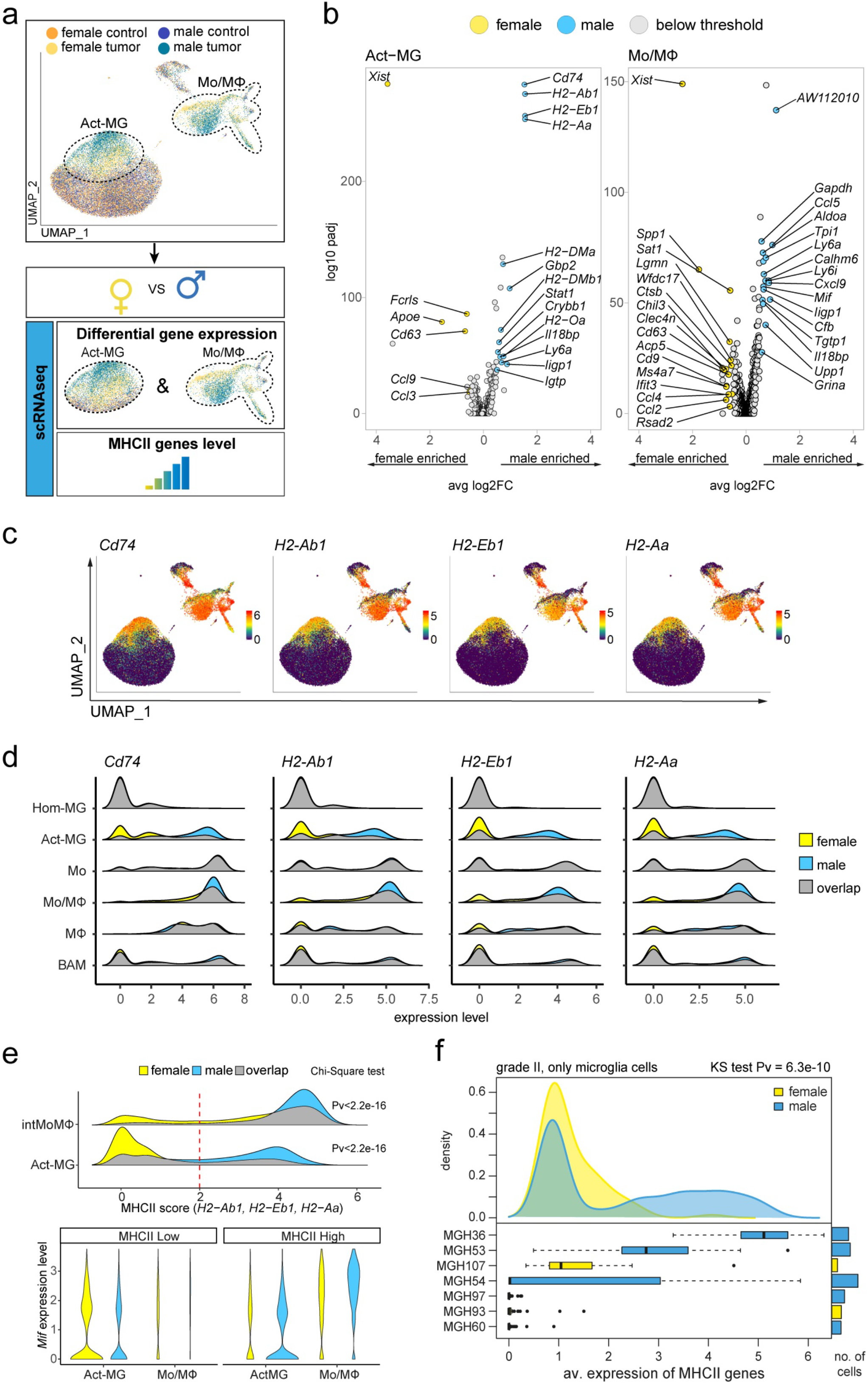
Expression *of* MHCII and *Cd74* genes is more abundant in microglia and monocytes/macrophages from gliomas in males. **a** Illustration of the analytical approach. UMAP plot demonstrates the distribution of male and female cells across cell clusters and reveals sex-enriched areas in Act-MG and Mo/MΦ. Differential gene expression analysis was performed for male vs female in Act-MG and Mo/MΦ groups, and expression level of top differentially expressed genes (DEG) verified across all cell groups. **b** Volcano plots depicting DEG across sexes in Act-MG and Mo/MΦ infiltrating gliomas. **c** Expression of the most highly upregulated genes from males. **d** Density plots show enrichment of male cells in MHCII genes- and *Cd74*-high expressing populations of Act-MG and intMoMΦ. **e** Violin and density plots demonstrate that *Mif* upregulation is limited to the intMoMΦ MHCII^hi^ cells. **f** Distribution of MHCII genes average expression in human single microglial cells. On the top yellow and blue distribution plots correspond to cells extracted from females and males respectively, and difference between the distribution was assessed with Kolmogorov-Smirnov test. Boxplot depicted below densities show distribution of the average MHCII complex genes expression in individual patients and bars on the right side correspond to the number of selected cells.

In Act-MG from males, among most highly upregulated genes are *H2-Ab1*, *H2-Eb1*, *H2-Aa* coding for the components of MHCII and *Cd74* – encoding an invariant MHCII chain implicated in folding and trafficking of the MHCII proteins **(Figure 5b)**. The increased expression of the MHCII genes and *Cd74* was found in the male-dominated cell clusters in the Act-MG, but also in the intMoMΦ **(Figure 5c)**. Accordingly, the cells with the high expression of MHCII and *Cd74* genes were enriched in Act-MG and intMoMΦ in males. This enrichment was not observed in Hom-MG, Mo, MΦ, and BAMs **(Figure 5d)**. IntMoMΦ in males upregulated *Mif* encoding a macrophage migration inhibitory factor. MIF stimulates CCL2-mediated macrophage migration and cell proliferation^53^, and in glioma TME it suppresses anti-tumoral microglial activity via activation of CD74^54^. The increased *Mif* expression in cells with high expression of MHCII genes was restricted to intMoMΦ and not detected in the Act-MG **(Figure 5e)**. Although high expression of MHCII genes was found in glioma-activated microglia and intMoMΦ from males, its implications in different subpopulations may differ in gliomas.

We performed analysis of the MHCII genes expression in microglia from WHO grade II glioma patients, using data from human single-cell studies^10,55^. In line with our findings on mouse gliomas, an average expression of the MHCII genes was higher in males **(Figure 5f)**. Female samples were underrepresented in this analysis despite including all data sets from single-cell studies on human gliomas where sex information was provided. In addition, we tested the human glioma expression data from The Cancer Genome Atlas (TCGA) to determine if sex has an impact on the expression of MHCII and *CD74* genes. Glioblastoma samples were not discriminated by expression of the selected MHCII and *CD74* genes (data not shown), irrespective of *IDH1* mutation status or macrophage content as estimated with the xCell^56^. However, the expression of MHCII and *CD74* genes stratified WHO grade II diffuse glioma patients into the female-enriched MHCII^lo^ and the male-enriched MHCII^hi^ groups **(Supplementary Figure 11)**.This observation shows that the differential regulation of genes coding for MHCII complex between sexes is not limited to a mouse glioma model, and those differences could be of clinical relevance.

## Discussion

In the present study, we have used flow cytometry and scRNA-seq to dissect the cellular and functional heterogeneity of GAMs. scRNA-seq of CD11b^+^ cells from naïve brains revealed considerable microglia heterogeneity suggesting existence of various functional states of microglia in the brain. Main clusters expressed homeostatic and signature genes (MG1)^31^, immediate early genes typical for transcriptionally active cells (MG2) or genes coding for signaling inhibitors and transcriptional repressors (MG3)^32,22^. Pre-MG cluster was enriched in the “microglia signature”, cytokines, chemokines and interferon response genes. Microglia and BMDMs accumulate in human glioblastomas and support glioma progression by augmenting tumor invasion, angiogenesis, and inducing immunosuppression^2^. Our main goal was to identify markers and functions of distinct myeloid subpopulations in murine malignant gliomas. Identifying specific roles of various subpopulations is critical for a cell population-specific intervention. Transcriptomic analyses of bulk CD11b^+^ infiltrates from human GBMs and murine gliomas showed a mixture of profiles characteristic for both pro- and antitumor phenotypes^4–6^. Cell separation based on CD45 expression has been criticized as CD45 can be upregulated in microglia under pathological conditions^57^. Herein, we show that *Tmem119* is highly expressed by microglia (both in control and glioma conditions), and *Lgals*3 (encoding Gal-3) by infiltrating monocytes/macrophages at RNA and protein levels. This pattern allows efficient separation of Tmem119^+^ and Gal-3^+^ cells within CD11b^+^ cells with flow cytometry. Interestingly, Tmem119 and Gal-3 separation of CD11b^+^ largely overlapped with CD45^hi/lo^ separation. Staining for Tmem119 and Gal-3 revealed a non-uniform cell distribution within the tumor, with predominance of monocytes/macrophages (Gal-3^+^) in the tumor core, and microglia (Tmem119^+^) occupying the tumor periphery. This observation is in agreement with results showing distinct spatial distribution of microglia and BMDMs in lineage tracing experiments ^48^, high resolution two photon imaging^58^, and a single-cell RNA-seq study on matched patient-derived samples from tumor core and periphery^9^.

Microglia and macrophage transcription regulatory networks adapt to changing environments^59,60^. Single cell RNA-seq studies on human GBMs suggested that microglia and monocytes/macrophages diminish their signature of origin, forming a phenotypic continuum and impeding clear separation^8,10^. In our study, the unsupervised cell clustering yielded three cell clusters representing microglia, monocytes/macrophages, and CNS BAMs. When we tested “microglia” and “macrophage” signatures, the “microglia signature” was indeed lower in glioma Act-MG, but its expression was still high and distinguishable from monocytes/macrophages. These observations demonstrate that the transcriptional signature of cell origin is retained in glioma TME.

Genetic lineage tracing studies showed BMDM accumulation in GL261 gliomas and transgenic RCAS-PDGF-B-HA gliomas, and distinct transcriptional networks associated with tumor-mediated education in microglia and recruited BMDMs^7,48^. We demonstrate that GL261 gliomas induce similar transcriptional networks in microglia and monocytes/macrophages, however the induction is stronger in monocytes/macrophages. This could be related to their prevalent localization within the tumor core, in contrast to microglia occupying tumor periphery^9,48^. Monocytes/macrophages express numerous genes related to immunosuppression, whereas, Act-MG show high expression of genes encoding chemokines, acting as chemoattractants from immune cells in mice^61^.

Monocytes/macrophages are heterogeneous and based on expression profiles we distinguished monocytes (*Ly6c2*^hi^*Ccr2*^hi^), intermediate state monocytes-macrophages (*Ly6c2*^hi^*Tgfbi*^hi^), and differentiated macrophages (*Ifitm2*^hi^, *S100a6*^hi^, *S100a11*^hi^) expressing a high level of *Cd274 (*coding for an immune checkpoint inhibitor - PD-L1). Those expression profiles together with patterns of cell distribution obtained with UMAP and FLE diffusion maps suggest that monocytes arrive as antitumor cells and undergo differentiation into pro-tumorigenic macrophages in the glioma microenvironment. We found only a partial overlap with the recently proposed macrophage-specific markers in gliomas^3,7,23^, but the proportion of monocyte/macrophage subpopulations may depend on the particular tumor stage. The occurrence of Mo/MΦ expressing an inflammatory *Il1b*, along with *Il1rn* and *Il18b* (coding for the inhibitors of pro-inflammatory cytokines) and *Cd274* is interesting for its clinical relevance, suggesting that pro-invasive and immunoregulatory functions are split between microglia and macrophages in TME, respectively.

Another important finding refers to sex-dependent differences in microglial responses to glioma. Bulk CD11b^+^ RNA-seq showed that microglia from male naïve mice express a higher level of MHCI and MHCII genes and are more reactive to ATP stimulation^13^. Estrogens mitigate inflammatory responses in microglia^62^, and female microglia have a higher neuroprotective capability^14^. We report that male microglia and intMoMΦ from the tumor-bearing hemispheres show higher expression of genes coding for MHCII components and Cd74. The analysis of TCGA and scRNA-seq human glioma datasets showed sex-related differences in MHCII complex and *CD74* genes in WHO grade II diffusive gliomas, where antitumor immunity operates and may influence outcomes. Such differences were not detected in highly immunosuppressed human GBMs. Although women are more susceptible to autoimmune diseases, men have a higher risk of death for a majority of malignant cancers^63^. In the immune checkpoint inhibitor therapy of various cancers, males presented better therapeutic outcomes^12^. While the source of sex differences in cancer incidence and outcome remains unknown, antitumor immunity is a plausible candidate.

In sum, microglia in a naïve brain are heterogeneous and represent various functional states. Glioma cells attract and polarize microglia and peripheral monocytes that migrate to CNS. Whereas infiltrating monocytes express some inflammatory markers, they likely differentiate into immunosuppressive macrophages within the tumor. Those cells retain their cell signatures, occupy different tumor niches, and display various degrees of glioma-induced activation and specific functions. Interestingly, we found the stronger upregulation of genes of the MCHII complex in microglia and fraction of monocytes/macrophages in male than female mice. Further studies on glioma immunopathology should explore this issue, ensure proper representation of both sexes and avoid extending findings from single-sex studies to the general population.

## Methods

### Animals

10-week-old male and female C57BL/6 mice were purchased from the Medical University of Bialystok, Poland. Animals were kept in individually ventilated cages, with free access to food and water, under a 12h/12h day and night cycle. All experimental procedures on animals were approved by the First Local Ethics Committee for Animal Experimentation in Warsaw (approval no 563/2018 and 764/2018).

### Implantation of GL261 luc^+^/tdT^+^ glioma cells

Mice (12-week-old) were kept under deep anesthesia with 2% isoflurane during surgery. Using a stereotactic apparatus, a single-cell suspension of GL261 luc^+^tdT^+^ cells (80 000 cells in 1μL of DMEM, Dulbecco modified essential medium) was implanted into the right striatum (+1 mm AP, −1.5 mm ML, −3 mm DV) at the rate of 0.25 μL per minute. In sham operated animals, 1 μL of DMEM was injected. To confirm the presence of the tumor, two weeks after implantation, animals received an intraperitoneal injection of 150 mg luciferin/kg body weight 10 min prior to imaging with the Xtreme *in vivo* bioluminescence imaging system [Bruker, Germany]. The images were acquired at medium binning with an exposure time of 2 min. X-ray images were acquired at the same mice position with the Xtreme equipment. The signal intensity of the region of interest (ROI) was computed using the provided software.

### Tissue dissociation

Two weeks after tumor implantation, mice with gliomas and naïve animals (controls) were perfused transcardially with cold phosphate-buffered saline (PBS) to clear away blood cells from the brain. Further processing was performed on the pooled tissue from 2 animals per replicate. The tumor-bearing hemispheres and corresponding hemispheres from naïve animals were dissociated enzymatically to obtain a single-cell suspension with a Neural Tissue Dissociation Kit with papain (Miltenyi Biotec) or 0.5 mg/mL DNase I (DN25, Sigma-Aldrich) in DMEM (Gibco, Germany) with 10% FBS for Tmem119 preparations and gentleMACS Octo Dissociator (Miltenyi Biotec) according to the manufacturer’s protocol. Next, the enzymatic reaction was stopped by the addition of Hank’s Balanced Salt Solution (HBSS) with calcium and magnesium (Gibco, Germany). The resulting cell suspension was filtered through a 70 μm and 40 μm strainer and centrifuged at 300g, 4 °C for 10 min. Next, myelin was removed by centrifugation on 22% Percoll gradient. Briefly, cells were suspended in 25 mL Percoll solution (18.9 mL gradient buffer containing 5.65 mM NaH_2_PO_4_H_2_O, 20 mM Na2HPO42(H2O), 135 mM NaCl, 5 mM KCl, 10 mM glucose, 7.4 pH; 5.5 mL Percoll (GE Healthcare, Germany); 0.6 mL 1.5 M NaCl), overlayered with 5 mL PBS and centrifuged for 20 min at 950 g and 4 °C, without acceleration and brakes. Next, cells were collected, washed with Stain Buffer (BD Pharmingen), quantified using an EVE™ Automatic Cell Counter (NanoEnTek Inc., USA), and split for CD11b+ FACS and cytometric analysis.

### Flow Cytometry

Samples were constantly handled on ice or at 4 °C avoiding direct light exposure. First, samples were incubated with LiveDead Fixable Violet Dead Cell Stain (ThermoFisher) or Fixable Viability Dye eF506 (eBioscience) in PBS for 10 min to exclude non-viable cells. Next, samples were incubated for 10 min with rat anti-mouse CD16/CD32 Fc Block™ (BD Pharmingen) in Stain Buffer (BD Pharmingen) to block FcγRIII/II and reduce unspecific antibody binding. Then, cell suspensions were incubated for 30 min with an antibody cocktail in Stain Buffer (BD Pharmingen). For cell surface antigens the following anti-mouse antibodies were used: from BD Pharmigen: CD45 (30-F11), CD11b (M1/70), Ly6C (AL-21); from BioLegend: CD49d (R1-2); from ThermoFisher: PD-L1 (MIH5); from Abcam: Tmem119 (106-6). For intracellular staining Foxp3/Transcription Factor Staining Buffer Set was used (eBioscence), following manufacturer’s instructions. For intracellular antigens the following anti-mouse Galectin 3 (M3/38 BioLegend) was used.).

Antibodies were titrated prior to staining to establish the amount yielding the best stain index. Samples were acquired using a BD LSR Fortessa Analyzer cytometer. Data were analyzed with FlowJo software (v. 10.5.3, FlowJo LLC, BD). Gates were set based on FMO (fluorescence minus one) controls and back-gating analysis. Percentages on cytograms were given as the percentage of a parental gate. All flow cytometry experiments were performed at the Laboratory of Cytometry, Nencki Institute of Experimental Biology. For reagent specifications, catalogue numbers and dilutions see the Supplementary Table 4.

### Sorting of CD11b^+^ cells by flow cytometry

Cells were incubated with LiveDead Fixable Violet Dead Cell Stain (ThermoFisher) in PBS for 10 min to exclude non-viable cells (Supplementary Figure 1d). Then, cells were suspended in Stain Buffer (BD Pharmingen) at a density of 1 million cells per 100 μL and stained with anti-mouse CD16/CD32 Fc Block™ (BD Pharmigen) for 10 min. Next, anti-mouse CD11b antibody (M1/70, BD Pharmigen) was added and cells were incubated for 20 min at 4 °C, washed with Stain Buffer, and sorted to 20% FBS in PBS.

### Single-cell RNA sequencing

Directly after sorting, cell quantity and viability of CD11b^+^ cells were measured, and a cell suspension volume equivalent to 5000 target cells was used for further processing. Preparation of gel beads in emulsion and libraries were performed with Chromium Controller and Single-Cell Gene Expression v2 Chemistry (or v3 for naïve vs sham-implanted experiment – Supplementary Figure 1) (10x Genomics) according to the Chromium Single-Cell 3’ Reagent Kits v2 (or v3) User Guide provided by the manufacturer. Libraries’ quality and quantity were verified with a High-Sensitivity DNA Kit (Agilent Technologies, USA) on a 2100 Bioanalyzer (Agilent Technologies, USA). Next, sequencing was run in the rapid run flow cell and paired-end sequenced (read 1 – 26 bp, read 2 – 100 bp) on a HiSeq 1500 (Illumina, San Diego, CA 92122 USA).

### Single-cell RNA-seq data preprocessing and normalization

Sequencing results were mapped to a mouse genome GRCm38 (mm10) acquired from the 10x Genomics website and quantified using a CellRanger v.3.0.1^64,65^ (https://support.10xgenomics.com/single-cell-gene-expression/software/pipelines/latest/installation). The total number of cells identified by the CellRanger was 41,059 (details in Supplementary Table 2). The median number of detected genes per cell was 1,059, and the median unique molecular identifiers (UMIs) per cell was 2,178. Data analysis was performed in R using Seurat v3^65,66^. Unless otherwise specified in the description, all other quantitative parameters were fixed to default values. To filter out possible empty droplets, low-quality cells, and possible multiplets, cells with <200 or >3,000 transcripts were excluded from the analysis. Additionally, cells of poor quality, recognized as cells with >5% of their transcripts coming from mitochondrial genes, were excluded from the downstream analysis. After applying these filters, 40,401 cells were present in the data set. Gene expression measurements for each cell were normalized by the total number of transcripts in the cell, multiplied by a default scale factor, and the normalized values were log-transformed (“LogNormalize” method). Following a Seurat workflow, for each replicate the 2,000 most highly variable genes were identified using variance stabilizing transformation (“vst”). To facilitate identification of cell types these gene sets were expanded by adding genes described as having important roles in immune cells (see Supplementary Table 3) and genes involved in cell cycle regulation^67^. This extension did not influence our conclusions.

### Identification of myeloid cells

Having two biological replicates for each sex and condition (female control, female tumor, male control, male tumor), data from corresponding samples were integrated using a Seurat v3 approach^65^. To avoid obtaining results fitted too closely to particular data set and therefore possibly failing to fit to additional data, firstly 2000 integration anchors (i.e., cells that are mutual nearest neighbors between replicates) were found. These anchors were then used as an input to the data sets integration procedure. Integrated data were scaled, and unwanted sources of variation, namely total number of counts per cell, percentage of transcripts coming from mitochondrial genes per cell, and cell cycle effect were regressed out, as described in a corresponding vignette [https://satijalab.org/seurat/v3.0/cell_cycle_vignette.html]. Data dimensionality reduction was performed using a principal component analysis (PCA), and the first 30 principal components were used in the downstream analyses. For each condition separately, the expression profiles were then clustered using an unsupervised, graph-based approach with the resolution parameter set to 0.3. Clustering results were visualized using two-dimensional t-Distributed Stochastic Neighbor Embedding (t-SNE)^68^. Based on expression of the reported/canonical markers, the clusters dominated by myeloid cells in four conditions were identified and further analyzed.

### Comparative analysis

The comparative analysis was based on the raw counts but limited to the previously selected profiles and genes (see above). For such a merged data set, a new set of the 2,000 most highly variable genes was identified using variance stabilizing transformation (“vst”), and this set was further expanded by adding the genes involved in cell cycle regulation. Computation of expression estimations, regression of the unwanted variation, and data dimensionality reduction were performed as described above. Next, the expression profiles were clustered using the same approach as above, but with a resolution parameter set to 0.6. After clustering, data were visualized using two-dimensional Uniform Manifold Approximation and Projection (UMAP)^69^. Based on expression of reported/canonical markers of myeloid cells, clusters with cells of interest (microglia, macrophages, and BAMs) were identified. Further analysis of the microglia cluster revealed that some sub-clusters cells originated in a significant majority from tumor samples. In contrast, there were no sub-clusters so strongly dominated by cells originated from control samples. Based on that observation, two subsets of microglial cells with distinct transcriptional profiles were identified: homeostatic microglia (Hom-MG) and activated microglia (Act-MG).

Differentially upregulated genes (signature genes) were found for each of the identity classes of interest. Significantly upregulated genes between compared groups were found using a Wilcoxon Rank Sum test implemented in Seurat v3 (min.pct = 0.25, only.pos = TRUE). These genes were subsequently used for the functional analysis and characterization of the identified clusters. Gene Ontology analysis was performed using the clusterProfiler package^70^.

### Human data analysis

MHCII plots were prepared using scRNA-seq glioma data from Tirosh *et al.* (2016) [GSE70630] and Venteicher *et al.* (2017) [GSE89567]. Only expression profiles from glioma WHO grade II were selected. For each patient top 10% cells with the highest microglia score were selected. For each of the selected cells an average expression of MHCII complex genes was computed [the list of genes in Table 3]. For TCGA data analysis the normalized expression values for low- and high-grade gliomas were downloaded from The Cancer Genome Atlas (TCGA) website (RNASeqV2 set available on 07/05/19). Sample annotations and *IDH1* mutation status were obtained from Ceccarelli and colleagues' study^71^. Content of immune cells was computed with xCell pre-calculated scores downloaded from the xCell website^56^. The genes encoding MHCII and CD74 proteins were selected based on the literature^17,72,73^. The expression profiles were clustered using hierarchical clustering. Significance of the clustering was computed using Fisher’s exact test. Separation of glioma samples was done for each grade separately.

### Immunohistochemistry on brain slices

For tissue collection for histology, mice were anesthetized and transcardially perfused with PBS and 4% paraformaldehyde (PFA). Brains were dissected and post fixed in 4% PFA overnight, then placed in 30% sucrose for 2 days, and then embedded in Tissue-Tek O.C.T Compound. Cryosections (10 μm) were cut and stored at −80°C. Cryosections were blocked in PBS containing 10% donkey serum in 0.1% Triton X-100 solution for 2 hours and incubated overnight at 4 °C with rat anti-Gal-3 and rabbit anti-Tmem119 antibody or rat anti-I-A/I-E (MHCII) and guinea pig anti-Tmem119 antibody. Next, sections were washed in PBS and incubated with corresponding secondary antibodies for 2 hours at room temperature. Nuclei were counter-stained with DAPI (0.1 mg/mL). Images were obtained on a Leica DM4000B fluorescent microscope. All antibodies were diluted in 0.1% Triton X-100/PBS solution containing 3% of donkey serum. For reagent specifications, catalogue numbers and concentrations see Supplementary Table 4.

## Data availability

Bam files and Seurat v3 processed gene expression matrix for each condition can be downloaded from the NIH GEO database (TBD).

## Code availability

The code to reproduce the analyses and figures described in this study can be found at: github.com/jakubmie/2019_OchockaSegit.

## Acknowledgments

We would like to thank Artur Wolny from Laboratory of Imaging Tissue Structure and Function at Nencki Institute of Experimental Biology for his help and assistance during analysis of the microscopic images, Kacper Łukasiewicz from University of California, Santa Cruz for his guidance in microscopy image processing in ImageJ software and Marcin Tabaka from Centre of New Technologies University of Warsaw for his guidance in scSVA analysis. Studies were supported by the National Science Centre, Poland (2017/27/B/NZ3/01605).

## Contributions

N.O., K.A.W., J.M. and B.K. conceived the study, N.O. and K.A.W. performed scRNA-seq experiments, P.S., J.M. and N.O. performed bioinformatical analysis, S.C. and J.S. performed cytometric experiments and analysis, K.W. performed immunohistochemical staining, N.O., P.S., J.M., S.C., K.W. prepared figures, J.M. and B.K. supervised the project, N.O., P.S., J.M. and B.K. wrote the manuscript. All authors have approved the submitted version.

