## Supplementary Information for "Single-cell RNA sequencing reveals functional heterogeneity and sex differences of glioma-associated brain macrophages"

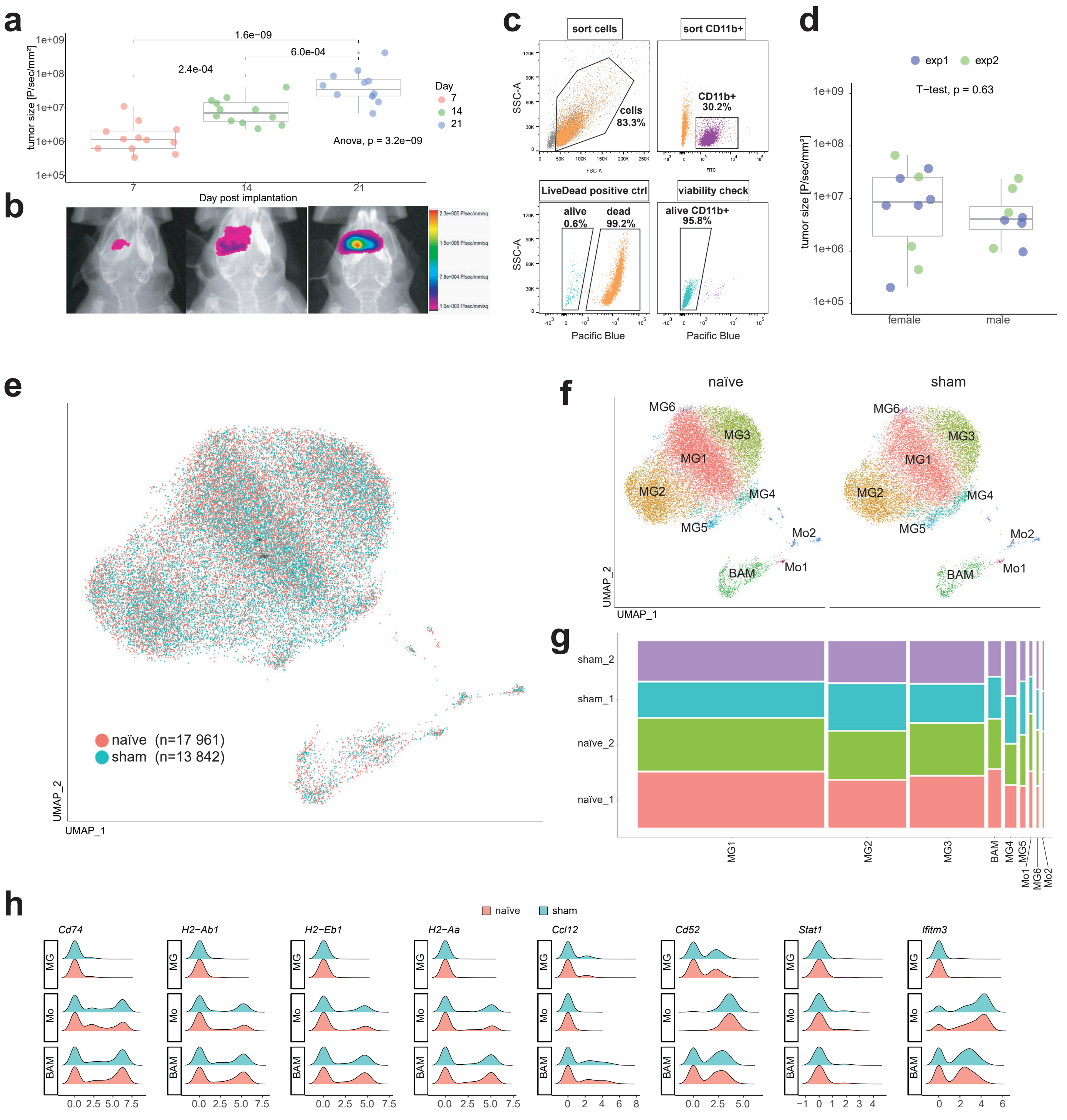


**Supplementary Figure 1.**
**(a)** Quantification of bioluminescence tumor imaging at 7, 14 and 21 day post-implantation. One-way ANOVA and Tukey's HSD post hoc test. **(b)** Representative tumor images (tumors for which the bioluminescent signal was closest to the median value in a given time point). **(c)** CD11b+ sorting strategy with labelling for live/dead cell showing high viability of sorted cells, with positive control for dead cells. **(d)** Quantification of bioluminescence tumor imaging male vs female animals for two biological replicates: exp1- animals sacrificed for scRNA-seq and cytometry measurements, exp2 – animals sacrificed for immunohistochemistry staining. **(e-h)** Single-cell RNA-seq for CD11b+ cells sorted from brains of naïve and sham-implanted animals. **(e)** UMAP shows uniform distribution of cells from naïve and sham-implanted animals. **(f)** UMAPs demonstrating that all obtained clusters were present for both naïve and sham condition **(g)** Proportion of cells from naïve and sham-implanted animals was comparable across all obtained clusters. **(h)** Expression level of genes showing upregulation in Act-MG compared to Hom-MG (Figure 4c), was not changed in sham-implanted compared to control animals.

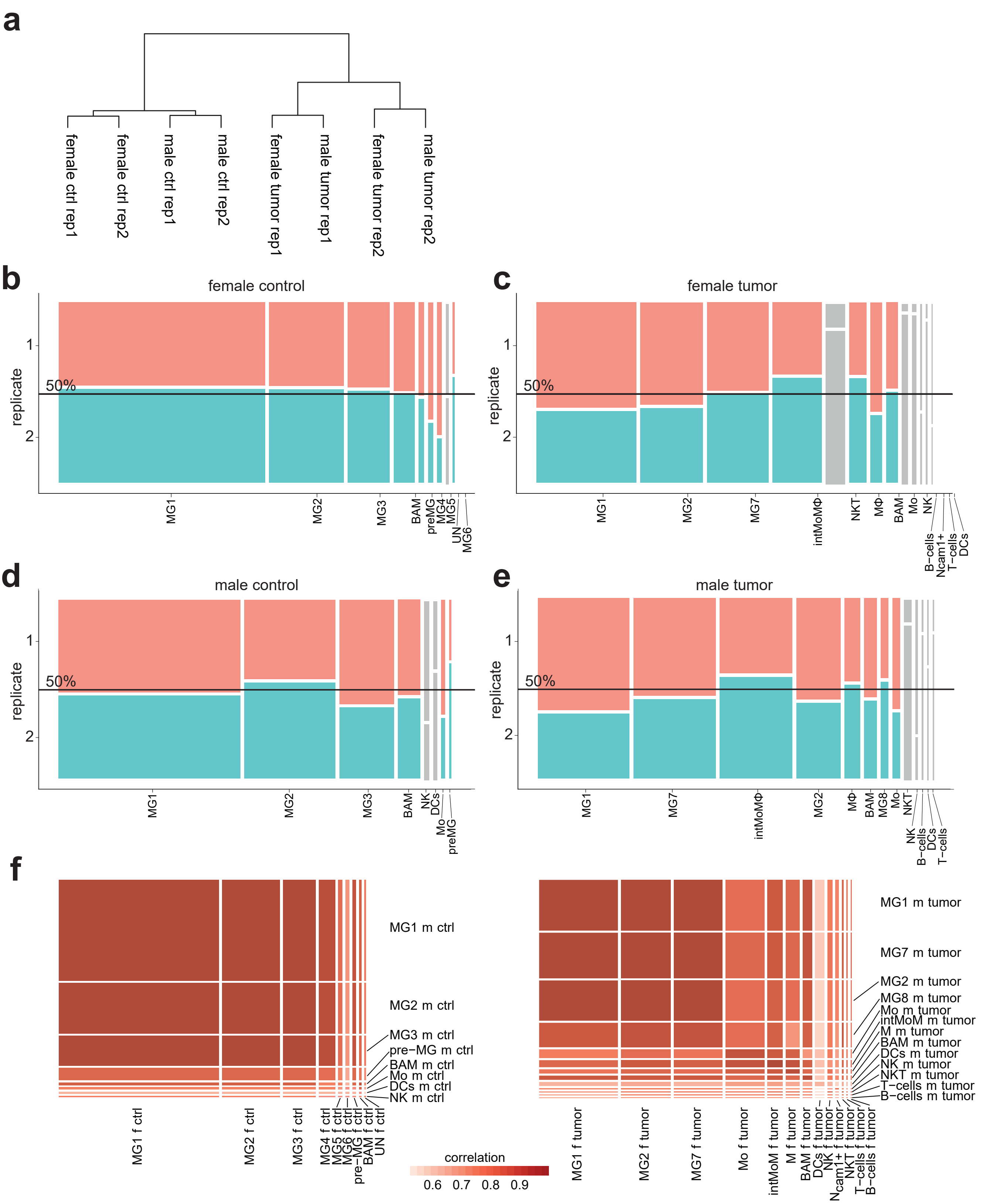


**Supplementary Figure 2.**
**(a)** Dendrogram showing results of unsupervised hierarchical clustering of pseudo-bulk gene expression profiles of each sample. **(b-e)** Percentage of cells from 2 replicates in clusters for **(b)** female control, **(c)** female tumor, **(d)** male control, **(e)** male tumor. Width of the each bar corresponds to the size of the cluster. Clusters that were selected for further analysis (Figure 2a) are in red and blue, the other clusters are in grey. **(f)** Correlation heatmaps comparing gene expression profiles of the identified clusters between sexes. Right and Left heatmap correspond to control and tumor samples respectively. Width and height of the cell represents fraction of cells joint into the corresponding cluster.

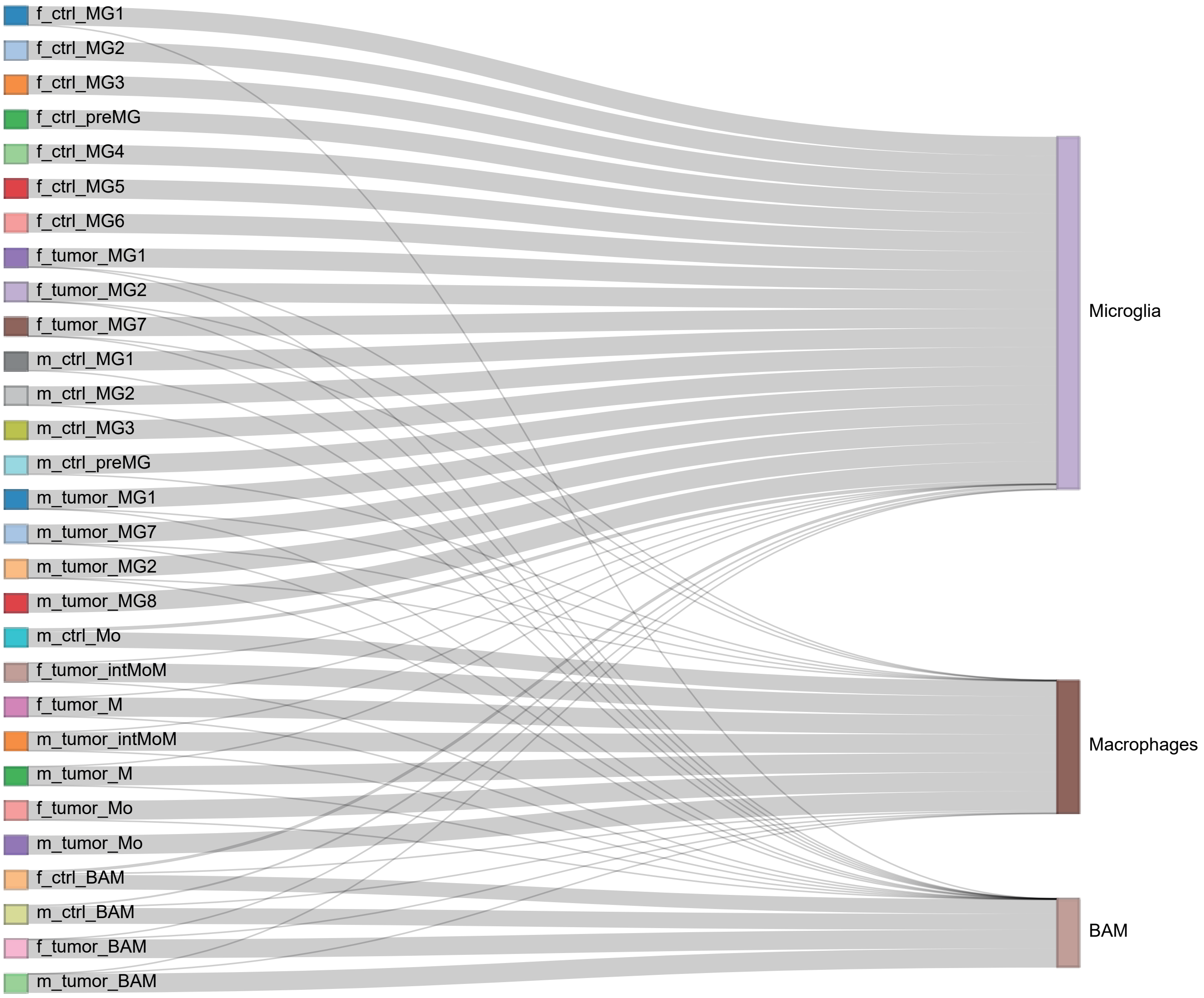


**Supplementary Figure 3.**
Flow diagram illustrating how cells from clusters obtained in analysis of each condition separately (Figure 1a) have transferred to clusters identified as Microglia, Macrophages, BAMs (Figure 2a), after merging selected cells from all samples into one dataset. Width of each link is proportional to the fraction of cells that have transferred from clusters on the left to clusters on the right side of the plot.

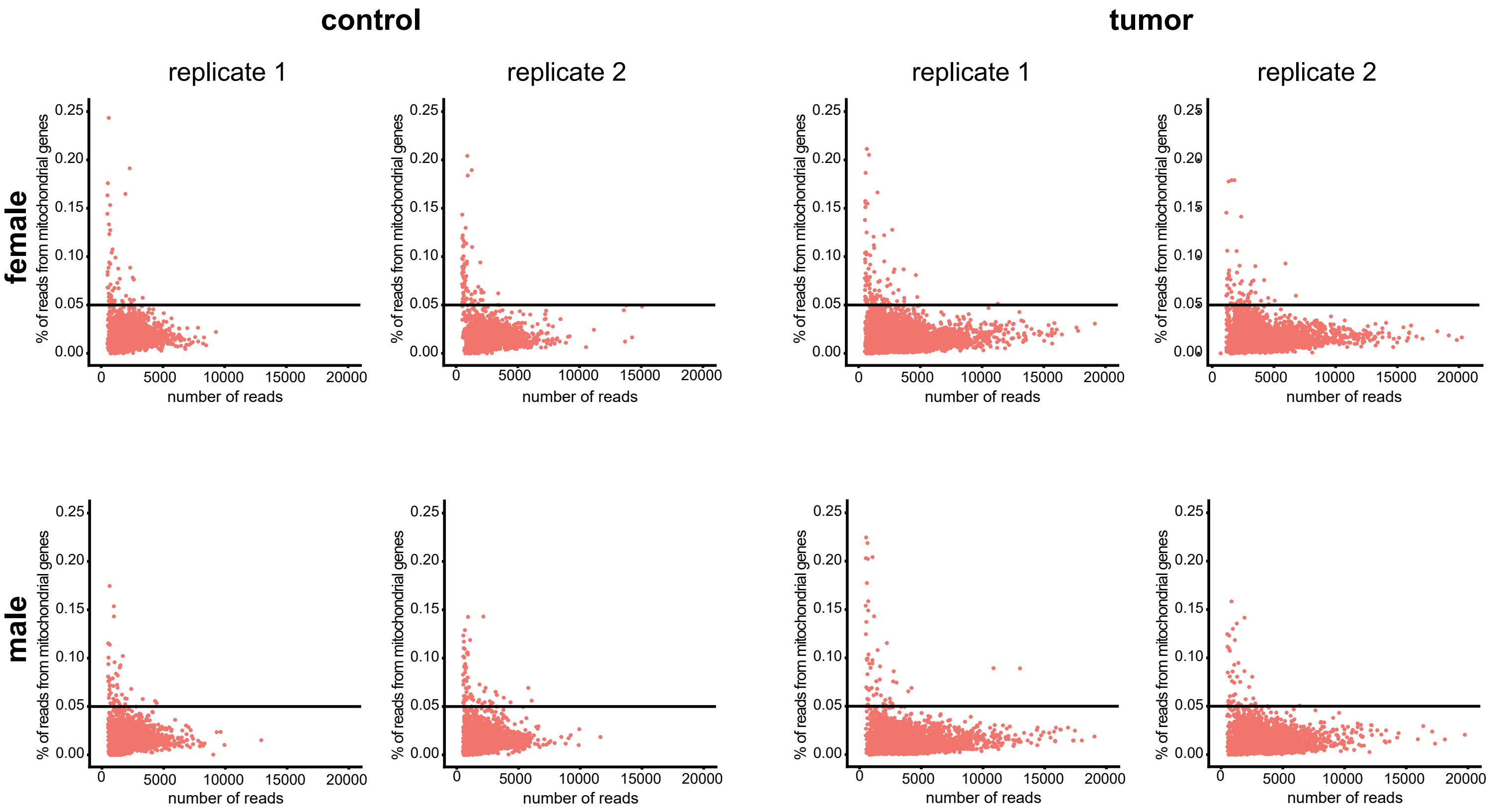


**Supplementary Figure 4.**
Scatter plots visualizing percentage of reads aligned to mitochondrial genes (Y axis) compared to the total number of reads (X axis). Each dot corresponds to individual cell. The figure is organized as Figure 1b.

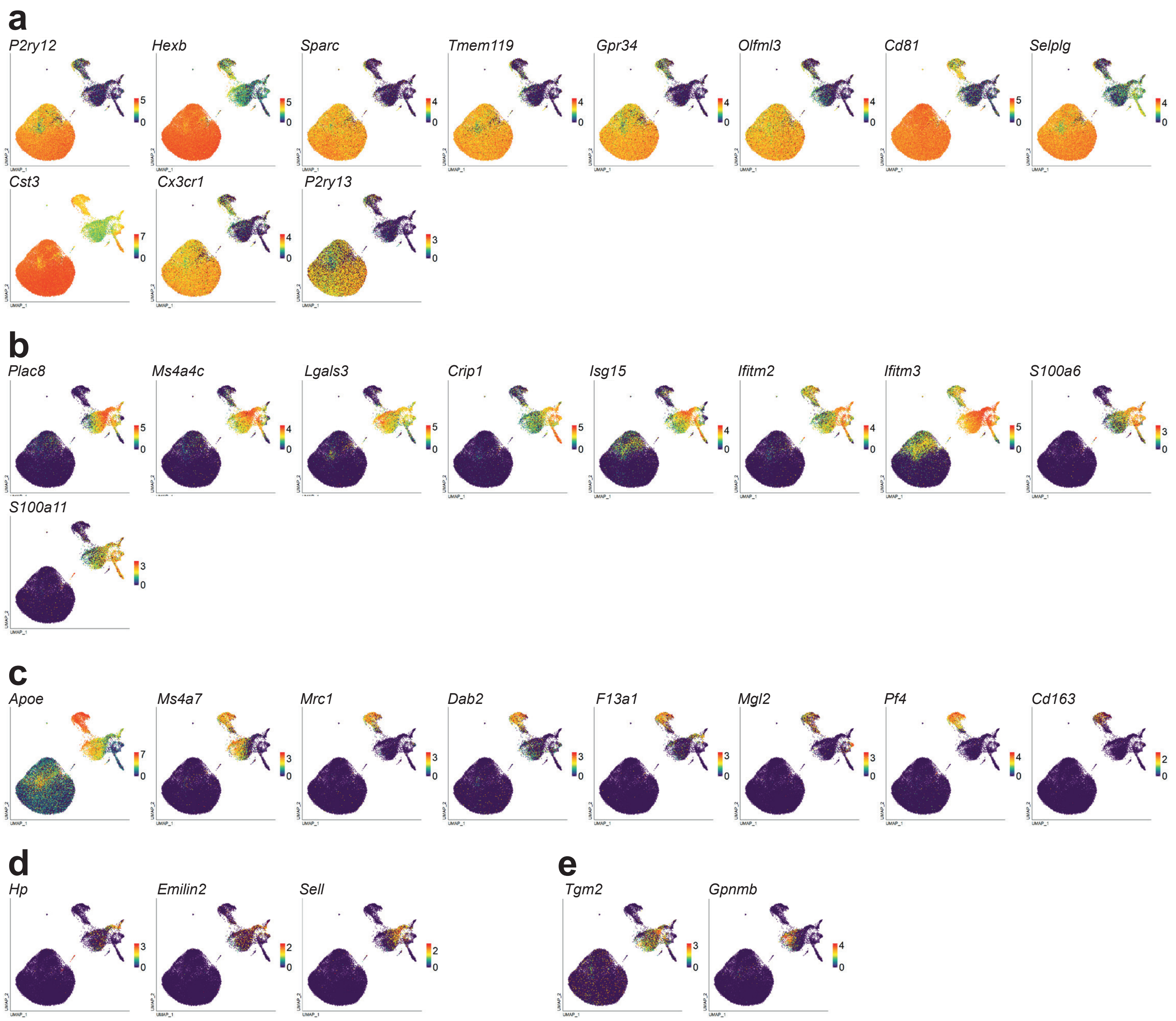


**Supplementary Figure 5.**
UMAP plots demonstrating the distribution of expression level for genes **(a)** highly expressed by microglia, **(b)** highly expressed by macrophages, **(c)** highly expressed by CNS Border Associated Macrophages (BAM), **(d)** proposed as markers of monocytes/macrophages infiltrating glioma TME by Haage et al. (2019), **(e)** proposed as markers of Glioma Associated Macrophages (GAM) by Walentynowicz et al. (2018).

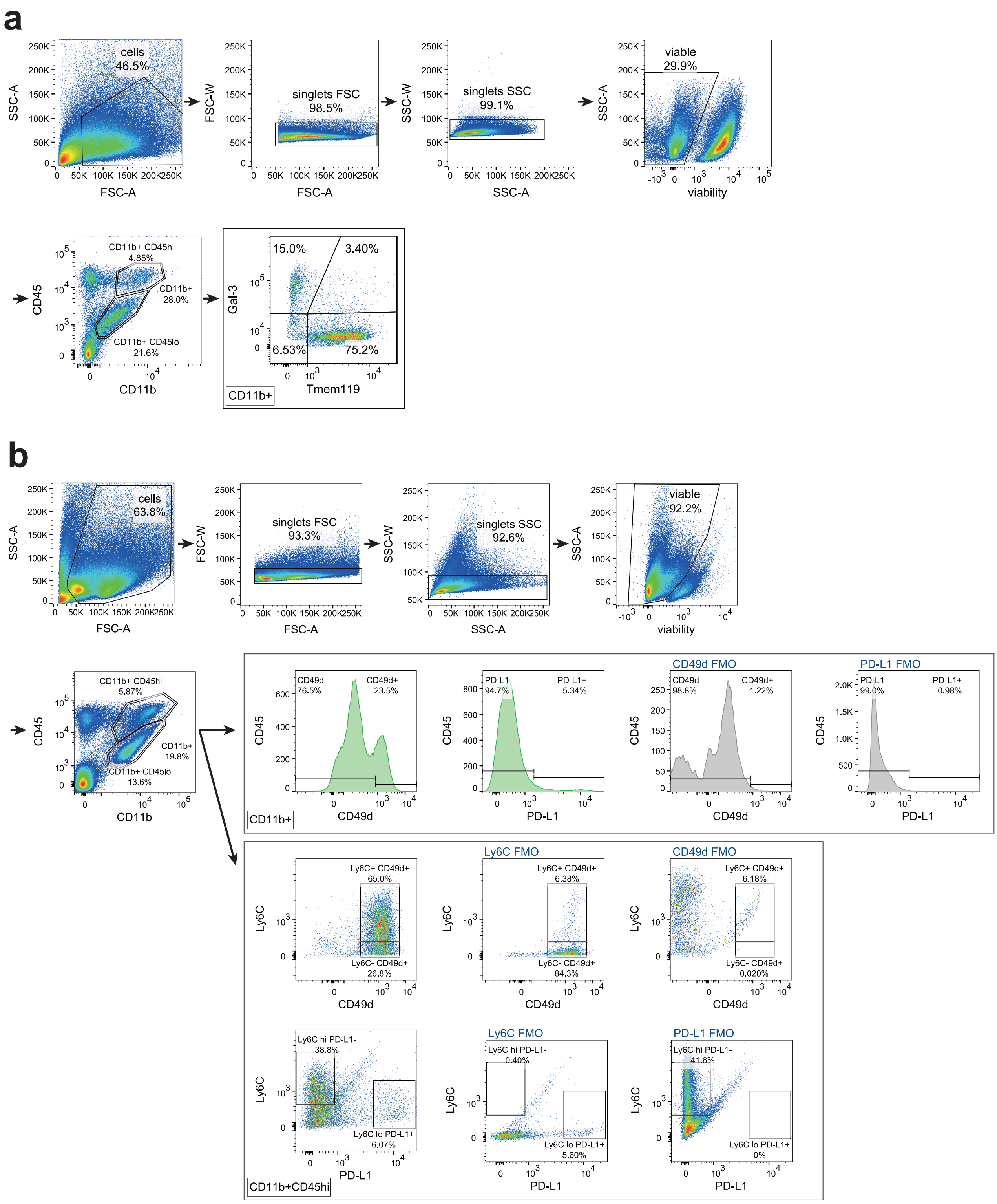


**Supplementary Figure 6.**

Gating strategy for flow cytometry analysis. **(a)** Gating for Tmem119 and Gal-3. **(b)** Gating for CD49d, PD-L1 and Ly6C. Events corresponding to cells were gated on SSC-A vs FSC-A plots, then doublets were excluded. Events in singlets gate were further analyzed for the uptake of Fixable Viability Dye (a) or LiveDead Violet dye (b) to exclude events corresponding to dead or damaged cells. For the gating of Tmem119+ and Gal-3+ events (a), gates were set on CD11b+ events. For CD49d+ and PD-L1+ events (b), gates were set on CD11b+ events, and for Ly6C vs CD49d and Ly6C vs PD-L1 analysis (b), gates were set on CD11b+CD45hi events. Gates were set based on backgating strategy or FMO controls. Tissues were dissociated enzymatically with DNase I (a) or papain-based enzyme mix (b) with simultaneous mechanical processing.

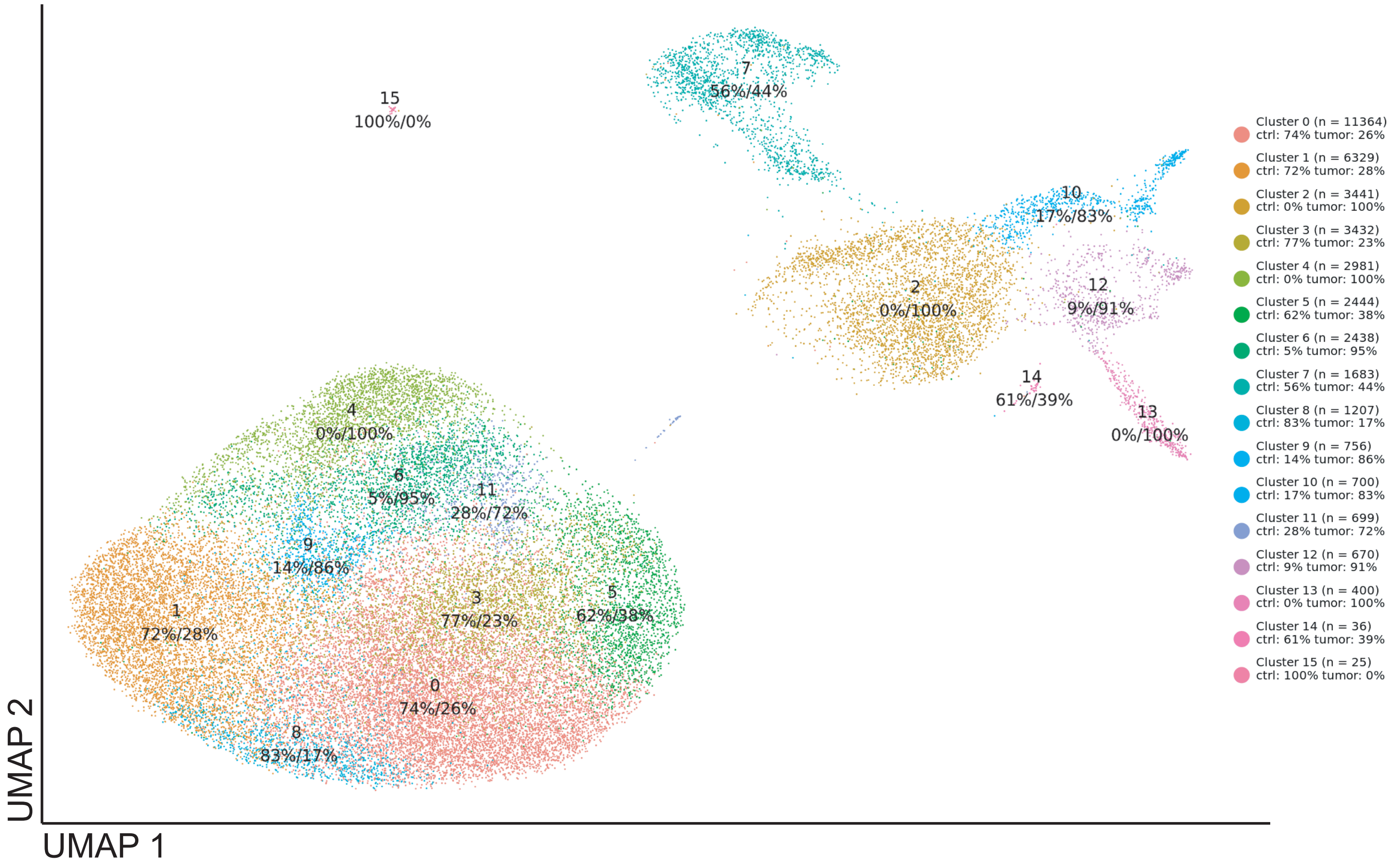


**Supplementary Figure 7.**
Results of unsupervised clustering of cells from three subpopulations of interest (corresponding to Figure 2a) from all 8 samples, visualized on UMAP plot. Clusters are marked with different colors and number of cells assigned to each cluster (n) is depicted in the legend. The shown percentages correspond to fractions of cells originating from control and tumor samples respectively.

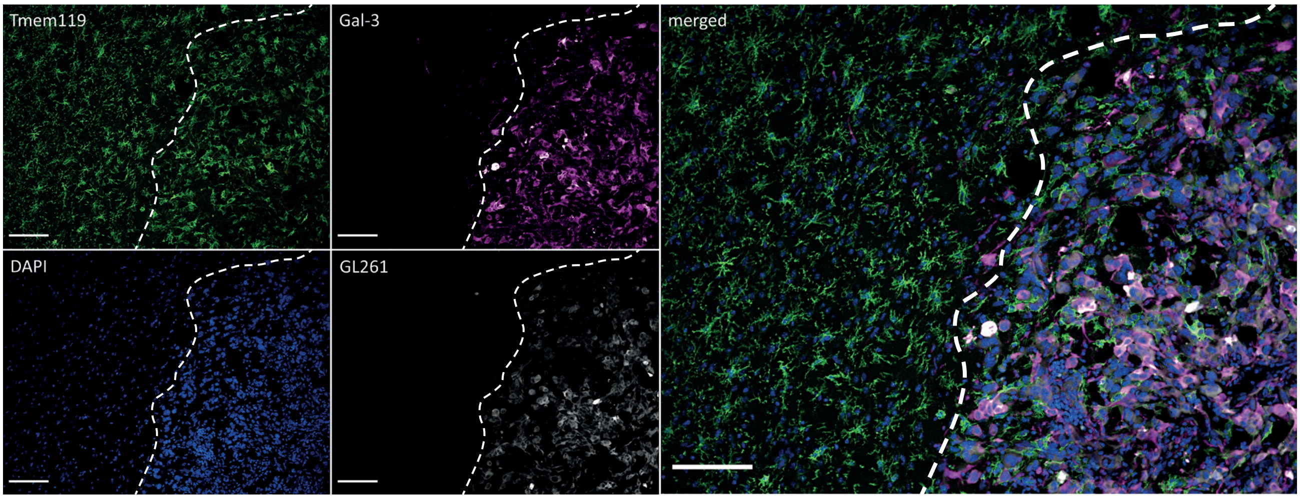


**Supplementary Figure 8.**
Immunohistochemical staining for microglia (Tmem119+) and Mo/MΦ (Gal-3+) shows the localization of specific immune cells within the tumor and its surroundings in male animal (for female see Figure 3d); a dashed line marks the tumor edge; scale – 100 µm.

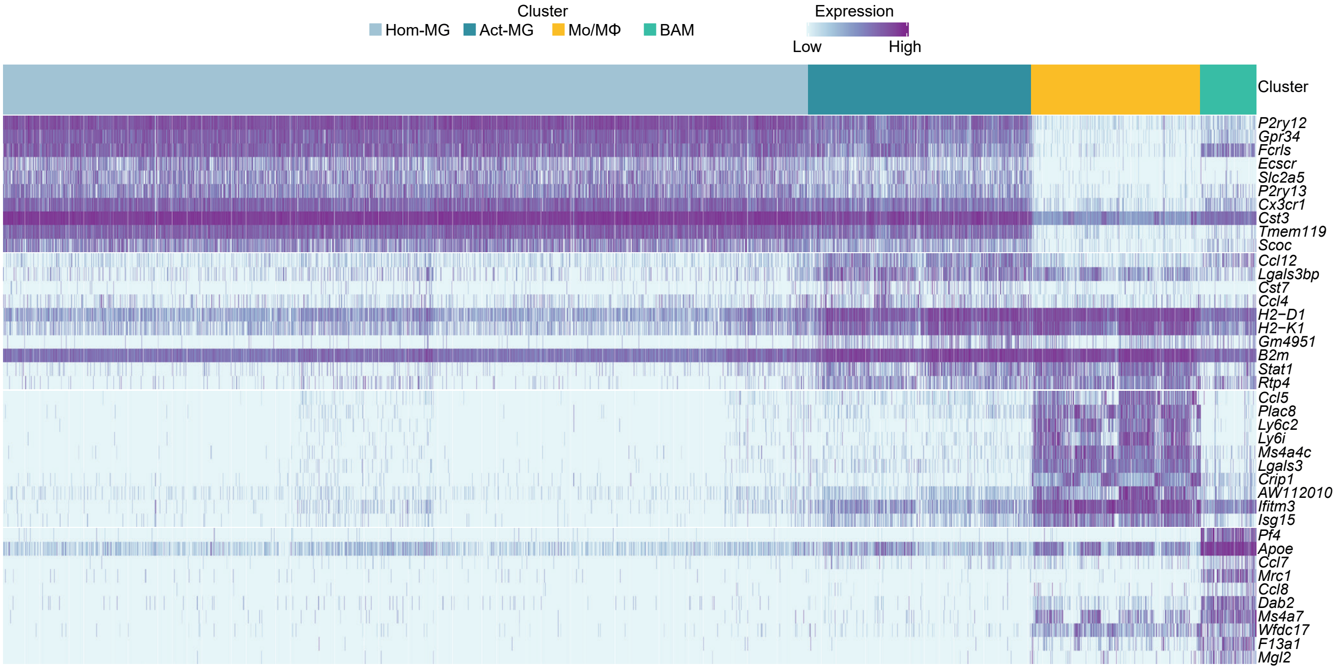


**Supplementary Figure 9.**
Heatmap demonstrating top 10 differentially expressed genes in Hom-MG, Act-MG, Mo/MΦ and BAM.

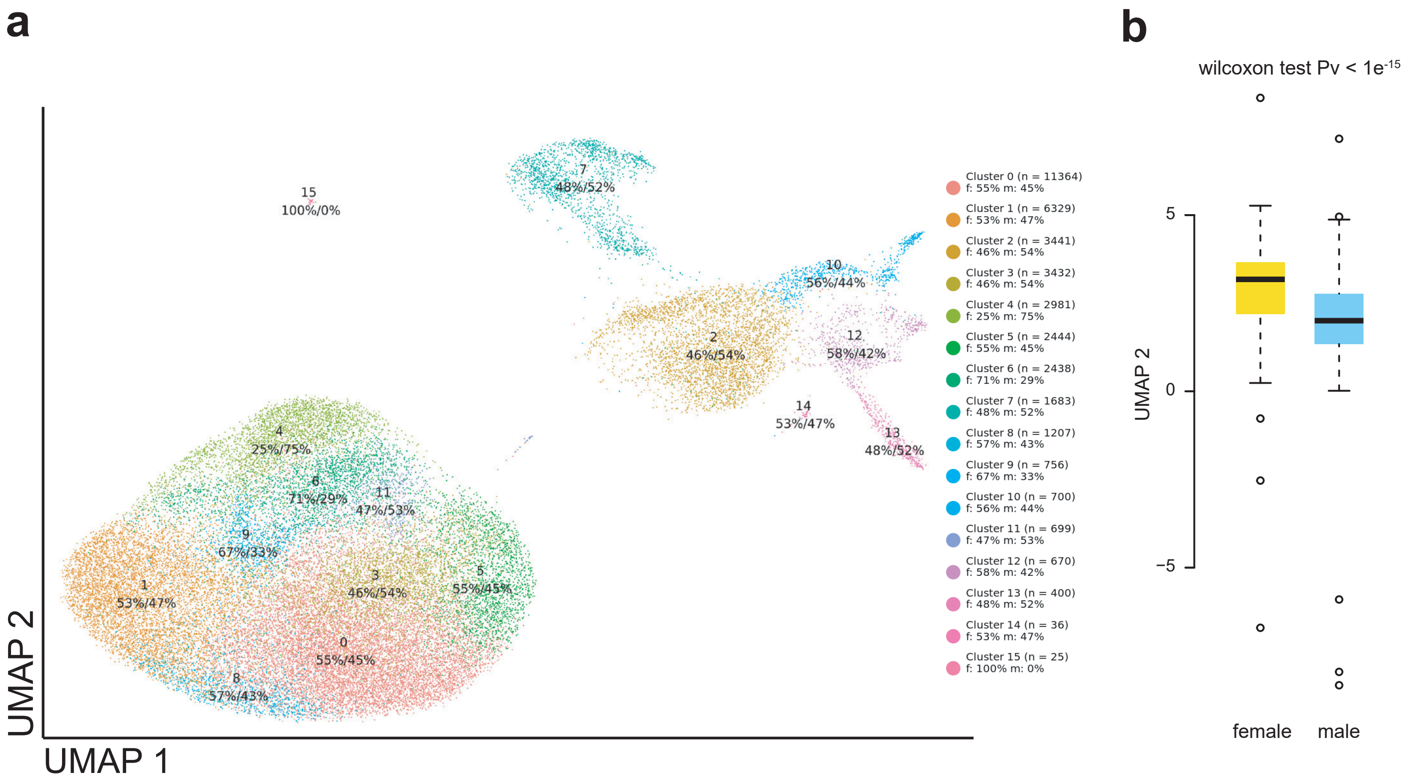


**Supplementary Figure 10.**
**(a)** Results of unsupervised clustering of cells from three subpopulations of interest (corresponding to Figure 2a) from all 8 samples, visualized on UMAP plot. Clusters are marked with different colors and number of cells assigned to each cluster (n) is depicted in the legend. The shown percentages correspond to fractions of cells originating from female and male samples respectively. **(b)** Comparison of UMAP 2 values for cells originating from female and male samples in cluster #2 identified as intermediate Monocyte-Macrophage cells - intMo/MΦ.

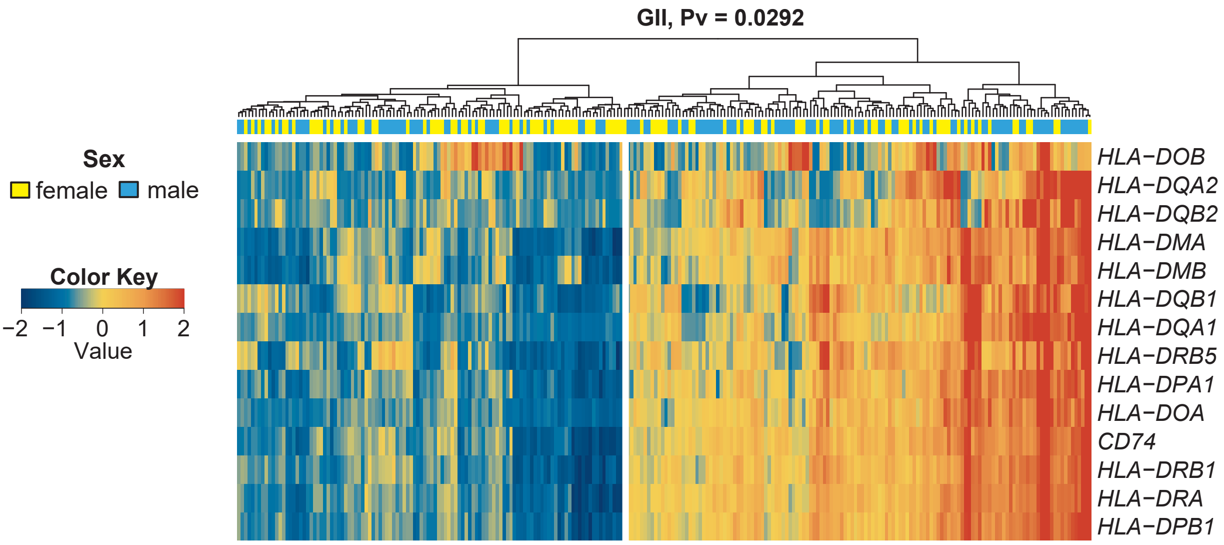


**Supplementary Figure 11.**
Normalized log2 RNA-seq counts for MHCII complex genes from TCGA WHO grade II glioma patients' data set shows significant differences between male and female glioma patients (Fisher's exact test).

**Supplementary Table 1.**

List of literature-based markers used to create an immune marker panel for characterizetion of cell identity of obtained clusters (Figure 1b,c).

| Gene | Target group | Ref. |
|---|---|---|
| Ptprc | hematopoietic cells | 1 |
| Itgam | myeloid cells | 1 |
| Cd14 | myelomonocytic cells | 1 |
| Tmem119 | microglia | 2,3,4 |
| Cx3cr1 | microglia | 2 |
| P2ry12 | microglia | 2,3 |
| P2ry13 | microglia | 2,3 |
| Gpr34 | microglia | 2,3 |
| Olfml3 | microglia | 2 |
| Selplg | microglia | 2,4 |
| Sparc | microglia | 2 |
| Fcrls | microglia | 2,3 |
| Siglech | microglia | 2 |
| Slc2a5 | microglia | 2,4 |
| Pf4 | microglia progenitors | 5 |
| F13a1 | microglia progenitors | 5 |
| Lyz2 | microglia progenitors | 5 |
| Ifit3 | microglia progenitors | 5 |
| Mcm5 | early microglia | 5 |
| Dab2 | early microglia | 5 |
| Cxcr2 | pre-microglia | 5 |
| Scd2 | pre-microglia | 5 |
| Psat1 | pre-microglia | 5 |
| Csf1 | pre-microglia | 5 |
| Crybb1 | pre-microglia | 5 |

| Gene | Target group | Ref. |
|---|---|---|
| Fcrls | pre-microglia | 5 |
| Selplg | adult microglia | 5 |
| Mafb | adult microglia | 5 |
| Pmepa1 | adult microglia | 5 |
| Cd14 | adult microglia | 5 |
| Lpl | disease associated microglia | 6 |
| Cst7 | disease associated microglia | 4,6 |
| Itga4 | macrophages | 7,9 |
| Tgfbi | macrophages | 8,9 |
| Ifitm2 | macrophages | 8,9 |
| Ifitm3 | macrophages | 8 |
| Tagln2 | macrophages | 8 |
| F13a1 | macrophages | 8 |
| Fpr3 | macrophages | 9 |
| Kynu | macrophages | 9 |
| S100a11 | macrophages | 9 |
| S100a6 | macrophages | |
| Tgm2 | Glioma Associated Microglia/Macrophages | 10 |
| Gpnmb | Glioma Associated Microglia/Macrophages | 10 |
| Emilin2 | macrophages in high-grade glioma enviorment | 11 |
| Gda | macrophages in high-grade glioma enviorment | 11 |
| Hp | macrophages in high-grade glioma enviorment | 11 |
| Sell | macrophages in high-grade glioma enviorment | 11 |
| Cd163 | Border Associated Macrophages | 12 |
| Mrc1 | Border Associated Macrophages | 12,13 |

| Gene | Target group | Ref. |
|---|---|---|
| Lyve1 | Border Associated Macrophages | 12 |
| Siglec1 | Border Associated Macrophages | 12 |
| Ly6c1 | monocytes | 14 |
| Ly6c2 | monocytes | 14 |
| Ccr2 | classical monocytes | 14,15 |
| Spn (CD43) | non-classical monocytes | 14 |
| Ly6g | Granulocytes | 16 |
| Cd24a | granulocytes/ dendritic cells | 16 |
| Itgax | dendritic cells | 17 |
| Bst2 | plasmocytoid dendritic cells | 17 |
| Ncam1 | NK cells | 18 |
| Klrb1c | NK cells | 18 |
| Klrk1 | NK cells | 18 |
| Ncr1 | NK cells | 18 |
| Cd2 | T-cells, NK cells | 1 |
| Cd3d | T cells | 1 |
| Cd3e | T cells | 1 |
| Cd3g | T cells | 1 |
| Cd4 | helper T cell | 1 |
| Cd8a | cytotoxic T cells | 1 |
| Cd8b1 | cytotoxic T cells | 1 |
| Cd19 | B-cells | 1 |
| Ms4a1 | B-cells | 1 |
| Sdc1 | B-cells | 1 |

**Supplementary Table 2.**
Number of identified cells, reads per cell and obtained stauration as well as analyzed cells and genes after filtration

| | female | | | | male | | | | | |
|---|---|---|---|---|---|---|---|---|---|---|
| | control rep1 | control rep2 | tumor rep1 | tumor rep2 | control rep1 | control rep2 | tumor rep1 | tumor rep2 | mean | sum |
| number of identified cells | 5 223 | 4 870 | 5 802 | 5 579 | 4 873 | 5 301 | 4 402 | 5 009 | 5 150 | 41 059 |
| number of reads per cell | 42 512 | 33 630 | 31 190 | 31 680 | 35 228 | 37 195 | 43 450 | 31 842 | 36 412 | |
| saturation | 90.1 | 87.0 | 83.8 | 84.7 | 88.6 | 89.5 | 86.2 | 84.9 | 87 | |
| number of cells remaining after filtration | 5 167 | 4 787 | 5 654 | 5 491 | 4 820 | 5 239 | 4 306 | 4 937 | 5 066 | 40 401 |
| number of genes remaining after filtration | 12 520 | 12 720 | 13 424 | 13 192 | 12 636 | 12 781 | 12 978 | 13 030 | | |

**Supplementary Table 3.**
The list of genes described as having important role in immune cells and involved in cell cycle regulation used to facilitate cell type identification.

| | | | | | | |
|---|---|---|---|---|---|---|
| *Anln* | *Cd27* | *Cxcl12* | *H2-Ea-ps* | *Itgb7* | *P2ry13* | *Timp2* |
| *Anp32e* | *Cd274* | *Cxcl16* | *H2-Eb1* | *Jun* | *Pcna* | *Tipin* |
| *Anxa1* | *Cd34* | *Cxcl2* | *H2-K1* | *Junb* | *Pf4* | *Tlr7* |
| *Anxa5* | *Cd38* | *Cxcl9* | *H2-Q7* | *Jund* | *Plac8* | *Tlr9* |
| *Apoe* | *Cd3d* | *Cxcr2* | *H2-T23* | *Kif11* | *Pmepa1* | *Tmem119* |
| *Arg1* | *Cd3e* | *Cybb* | *H2afv* | *Kif20b* | *Pola1* | *Tmem123* |
| *Ass1* | *Cd3g* | *Cycs* | *H2afy* | *Kif23* | *Pold3* | *Tmpo* |
| *Atad2* | *Cd4* | *Dab2* | *H2afy2* | *Kif2c* | *Prim1* | *Tnf* |
| *Aurka* | *Cd40* | *Dlgap5* | *H2afz* | *Kit* | *Pros1* | *Top2a* |
| *Aurkb* | *Cd52* | *Dscc1* | *H3f3a* | *Klf2* | *Psat1* | *Tpx2* |
| *Basp1* | *Cd63* | *Dtl* | *H3f3b* | *Klf4* | *Psrc1* | *Trafd1* |
| *Bhlhe41* | *Cd72* | *Dusp1* | *Hells* | *Klf6* | *Ptprc* | *Trem2* |
| *Birc5* | *Cd74* | *Dusp2* | *Hjurp* | *Klrb1c* | *Rad51* | *Ttk* |
| *Blm* | *Cd81* | *Dusp3* | *Hmgb2* | *Klrk1* | *Rad51ap1* | *Tubb4b* |
| *Bmp2k* | *Cd8a* | *Dusp5* | *Hmmr* | *Lbr* | *Rangap1* | *Tyms* |
| *Brip1* | *Cd8b1* | *Dusp6* | *Hp* | *Lgals3bp* | *Relb* | *Ube2c* |
| *Bst2* | *Cd9* | *E2f8* | *Ier5* | *Lpl* | *Rfc2* | *Ubr7* |
| *Bub1* | *Cdc20* | *Ecscr* | *Ifi205* | *Ly6c1* | *Rnh1* | *Uhrf1* |
| *Casp8ap2* | *Cdc25c* | *Ect2* | *Ifit3* | *Ly6c2* | *Rpa2* | *Ung* |
| *Cbx5* | *Cdc45* | *Egr1* | *Ifitm2* | *Ly6g* | *Rrm1* | *Usp1* |
| *Ccl12* | *Cdc6* | *Emilin2* | *Ifitm3* | *Ly6i* | *Rrm2* | *Wdr76* |
| *Ccl17* | *Cdca2* | *Exo1* | *Il10* | *Lyz1* | *S100a11* | *Xaf1* |
| *Ccl2* | *Cdca3* | *F13a1* | *Il10rb* | *Mafb* | *S100a4* | *Zbp1* |
| *Ccl22* | *Cdca7* | *Fcer1a* | *Il12a* | *Mcm2* | *S100a6* | *Zfhx3* |
| *Ccl24* | *Cdca8* | *Fcrls* | *Il12b* | *Mcm4* | *Sall1* | *Zfp691* |
| *Ccl3* | *Cdk1* | *Fen1* | *Il13ra1* | *Mcm5* | *Samhd1* | |
| *Ccl4* | *Ceacam1* | *Fgr* | *Il15* | *Mcm6* | *Scd2* | |
| *Ccl5* | *Ceacam3* | *Flt3* | *Il17ra* | *Mef2c* | *Sdc1* | |
| *Ccl6* | *Cebpa* | *Fos* | *Il18* | *Mki67* | *Sell* | |
| *Ccl7* | *Cebpd* | *Fosb* | *Il18bp* | *Mndal* | *Selplg* | |
| *Ccl8* | *Cenpa* | *G2e3* | *Il1a* | *Mrc1* | *Siglecf* | |
| *Ccl9* | *Cenpe* | *Gas2l3* | *Il1b* | *Ms4a1* | *Siglech* | |
| *Ccnb2* | *Cenpf* | *Gas6* | *Il1rn* | *Msh2* | *Slamf7* | |
| *Ccne2* | *Chaf1b* | *Gbp2* | *Il2* | *Nampt* | *Slbp* | |
| *Ccr1* | *Ckap2* | *Gbp5* | *Il2rb* | *Nasp* | *Slc2a5* | |
| *Ccr2* | *Ckap2l* | *Gda* | *Il2rg* | *Ncam1* | *Slc39a1* | |
| *Ccr3* | *Ckap5* | *Gins2* | *Il3ra* | *Ncapd2* | *Smc4* | |
| *Ccr7* | *Cks1b* | *Gmnn* | *Il4i1* | *Ncoa3* | *Socs1* | |
| *Ccrl2* | *Cks2* | *Gpnmb* | *Il5ra* | *Ncr1* | *Socs2* | |
| *Cd14* | *Clspn* | *Gpr141* | *Il6* | *Ndc80* | *Socs3* | |
| *Cd163* | *Crybb1* | *Gpr34* | *Il6ra* | *Nek2* | *Sparc* | |
| *Cd180* | *Csf1* | *Gtse1* | *Irf7* | *Nfia* | *Spp1* | |
| *Cd19* | *Cst3* | *H1f0* | *Irf8* | *Nfkbia* | *Tacc3* | |
| *Cd2* | *Cst7* | *H1fx* | *Itga2* | *Notch1* | *Tagln2* | |
| *Cd200* | *Cstb* | *H2-Aa* | *Itga2b* | *Npc2* | *Tgfbi* | |
| *Cd200r4* | *Ctcf* | *H2-Ab1* | *Itga4* | *Nuf2* | *Tgfbr1* | |
| *Cd209a* | *Cx3cr1* | *H2-D1* | *Itgam* | *Nusap1* | *Tgm2* | |
| *Cd24a* | *Cxcl10* | *H2-DMa* | *Itgax* | *P2ry12* | *Thy1* | |

**Supplementary Table 4.**
Specifications, catalog numbers and dilutions of reagents used for immunohistochemistry and flow cytometry.

| Reagent | Manufacturer | Cat. number | Clone | Fluorophore | Dilution |
|---|---|---|---|---|---|
| LiveDead Fixable Violet Dead Cell Stain | ThermoFisher | L34955 | - | - | 1:1000 |
| Fixable Viability Dye eF506 | eBioscience | 65-0866 | - | - | 1:1000 |
| Stain Buffer | BD Pharmingen | 554656 | - | - | - |
| Foxp3 Transcription Factor Staining Buffer | eBioscience | 00-5523-00 | - | - | - |
| anti-mouse CD16/CD32 Fc Block | BD Pharmingen | 553141 | - | - | 1:250 |
| anti-CD45 | BD Pharmingen | 561868 | 30-F11 | PE-Cy7 | 1:800 |
| anti-CD11b | BD Pharmingen | 557960 | M1/70 | Alexa Fluor 700 (flow cytometry) | 1:800 |
| anti-CD11b | BD Pharmingen | 553310 | M1/70 | FITC (FACS) | 1:800 |
| anti-Ly6C | BD Pharmingen | 560525 | AL-21 | PerCP-Cy5.5 | 1:100 |
| anti-CD49d | BioLegend | 103605 | R1-2 | FITC | 1:400 |
| anti-PD-L1 | ThermoFisher | 63-5982-82 | MIH5 | SuperBright600 | 1:100 |
| anti-Tmem119 | Abcam | ab210405 | 106-6 | unconjugated (rabbit) | 1:400 |
| anti-rabbit Alexa Fluor 488 pAb | Abcam | ab150077 | - | Alexa Fluor 488 | 1:1000 |
| anti-Gal-3 | eBioscience | 125408 | M3/38 | Alexa Fluor 648 | 1:200 FC<br>1:100 IF |
| anti-TMEM119 pAb | Synaptic Systems | 400002 | - | - | 1:500 |
| anti-MHC II | ThermoFisher | 14-5321-82 | M5/114.15.2 | - | 1:200 |
| anti-TMEM119 pAb | Synaptic Systems | 400004 | - | - | 1:500 |
| anti-rabbit Alexa Fluor 488 pAb | Invitrogen | A21206 | - | Alexa Fluor 488 | 1:1000 |
| anti-rat Alexa Fluor pAb | Invitrogen | A21208 | - | Alexa Fluor 488 | 1:1000 |
| anti-guinea pig Cy5 pAb | Jackson Immunoresearch | 706-175-148 | - | Cy5 | 1:1000 |
